# The structural unit of melanin in the cell wall of the fungal pathogen *Cryptococcus neoformans*

**DOI:** 10.1101/644484

**Authors:** Emma Camacho, Raghav Vij, Christine Chrissian, Rafael Prados-Rosales, David Gil, Robert N. O’Meally, Radames J.B. Cordero, Robert N. Cole, J. Michael McCaffery, Ruth E. Stark, Arturo Casadevall

## Abstract

Melanins are synthesized macromolecules that are found in all biological kingdoms. These pigments have a myriad of roles that range from microbial virulence to key components of the innate immune response in invertebrates. Melanins also exhibit unique properties with potential applications in physics and material sciences, ranging from electrical batteries to novel therapeutics. In the fungi, melanins such as eumelanins, are components of the cell wall that provide protection against biotic and abiotic elements. Elucidation of the smallest fungal cell wall-asociated melanin unit that serves as a building block is critical to understand the architecture of these polymers, its interaction with surrounding components, and their functional versatility. In this study, we used isopycnic gradient sedimentation, NMR, EPR, high-resolution microscopy, and proteomics to analyze the melanin in the cell wall of the human pathogenic fungus *Cryptococcus neoformans*. We observed that melanin is assembled into the cryptococcal cell wall in spherical structures of ∼200 nm in diameter, termed melanin granules, which are in turn composed of nanospheres of ∼30 nm in diameter, the fungal melanosomes. We noted that melanin granules are closely associated with proteins that may play critical roles in the fungal melanogenesis and the supramolecular structure of this polymer. Using this structural information, we propose a model for *C. neoformans* melanization that is similar to the process used in animal melanization and is consistent with the phylogenetic relatedness of the fungal and animal kingdoms.

## INTRODUCTION

Melanins (Greek melano, meaning dark or black) are perhaps the most enigmatic biopolymers in the biosphere despite being ubiquitous in nature (1). These pigments carry out a vast array of functions in all biological kingdoms including thermoregulation, radical scavenging, energy transduction, camouflage, invertebrate immunity and sexual display (2–4). In the microbial world melanins contribute to the virulence of many pathogenic microbes. Melanins are produced through the oxidation and polymerization of phenolic/indolic precursors; notably, these polymers are characterized by a strong negative charge, high molecular weight, and hydrophobic nature (5). In the fungal field, black-to-brown insoluble pigments designated as eumelanins confer greater fitness to melanotic species *in vivo*, thereby contributing to their virulence (6). In *Cryptococcus neoformans*, a facultative intracellular pathogen, melanized cells are able to modulate the immune response of the host through multiple mechanisms e.g. altering cytokine profiles (7), reducing antibody-mediated phagocytosis, and decreasing the toxicity of microbial peptides, reactive oxygen species (ROS), and antifungal drugs (8,9). Recent synthesis of a soluble melanin (homogentesic acid and L-DOPA at a 1:1 ratio), demonstrated that it suppresses the production of cytokines and ROS under stimulation of fungal components thus verifying melanin’s ability to modulate the immune system (10).

*C. neoformans* is a unique system for the study of melanin biology because unlike other melanotic fungi, only exogenously provided catecholamines or diphenolic substrates, such as dihydroxyphenylalanine (L-DOPA), are utilized for melanin synthesis and not as an energy source (11–13). Consequently, it is possible to selectively label the melanin polymer with NMR- or radioactive isotopes, a property that has been exploited for melanin structural studies (14–19). Melanization of *C. neoformans* results in the deposition of the polymer at the innermost surface of the cell wall, which becomes filled throughout over time (20). The pigment is composed of closely packed spherical particles ranging from 40 to 130 nm in diameter, which are found arranged in concentric layers in the cell wall (21). Early ultrastructural observations in *Agaricus bisporus (22), Fonsecaea pedrosoi* (23), *Cladosporium carrionii* and *Hormoconis resinae* (24) suggested compartmentation of fungal melanins into organelles akin to mammalian melanosomes (25). Evidence for trans-cell-wall vesicular transport in fungi associated with virulence (26), supported the hypothesis that fungal melanization may occur within specialized vesicles (27–30).

Studies using isotopically-labeled precursors in conjunction with high resolution solid-state nuclear magnetic resonance (ssNMR), revealed that the melanin polymer is likely to be covalently bonded to cell-wall chitin and also found strongly associated with other non-pigment cellular moieties including polysaccharides such as chitosan and vesicle and/or plasma membrane derived lipids (16,19). These components serve as the scaffold for melanin synthesis. Indeed, *C. neoformans* strains with aberrant chitin and/or chitosan biosynthesis are unable to retain the pigment within the cell wall and display a leaky-melanin phenotype (31–34), whereas boosting cell-wall chitin/chitosan content increases melanin deposition and assembly (14). The association between fungal melanin and chitin was initially reported in *Aspergillus nidulans* (35), followed by work done on *Exophilia dermatitides* (36) and *Candida albicans* (29). The concept of a fibrillary matrix upon which melanin is deposited extends to other species including human eumelanin, where its known to also play a critical role in absorbing toxic oxidative melanin intermediates and provide the elliptical shape to melanosomes (37).

In addition to the relevance of melanin to all biota and its great impact on fungal pathogenesis, this polymer has numerous potential applications in biophysics, material sciences, and even in the cosmetics and health care industries (38–40). The remarkable potential of melanin derives both from its chemical composition on the molecular level and its versatility as a component of the supramolecular structure within a cellular milieu. Investigation of the melanin structure is a notoriously challenging task due to the insoluble, amorphous, and heterogeneous nature of this architecturally complex polymer (41). Despite these challenges, over the past two decades a wealth of molecular level structural information has been uncovered using multiple non-destructive methodologies among them: X-ray scattering, ssNMR, electron microscopy, and atomic force microscopy (16,42–46). Current views of eumelanin molecular architecture posit that these pigments are composed of a common chemical motif and arranged as graphite-like *π*-stacked planar sheets of locally ordered monomers, mainly derivatives of 5,6-dihydroxyindole (DHI) (47,48), which give rise to subunit nanoparticles that then undergo spontaneous aggregation until submicron particles are formed (45,46). Nonetheless, natural eumelanins differ in structure from synthetic melanin and neuromelanin (44,45), suggesting that the mode of synthesis can affect the final structure. A graphical schematic highlighting the current views on eumelanin synthesis and the relationship of melanin to the fungal cell wall, capsule and cell body shows that melanin resides in the cell wall (**Figure 1**).

**Fig. 1.**
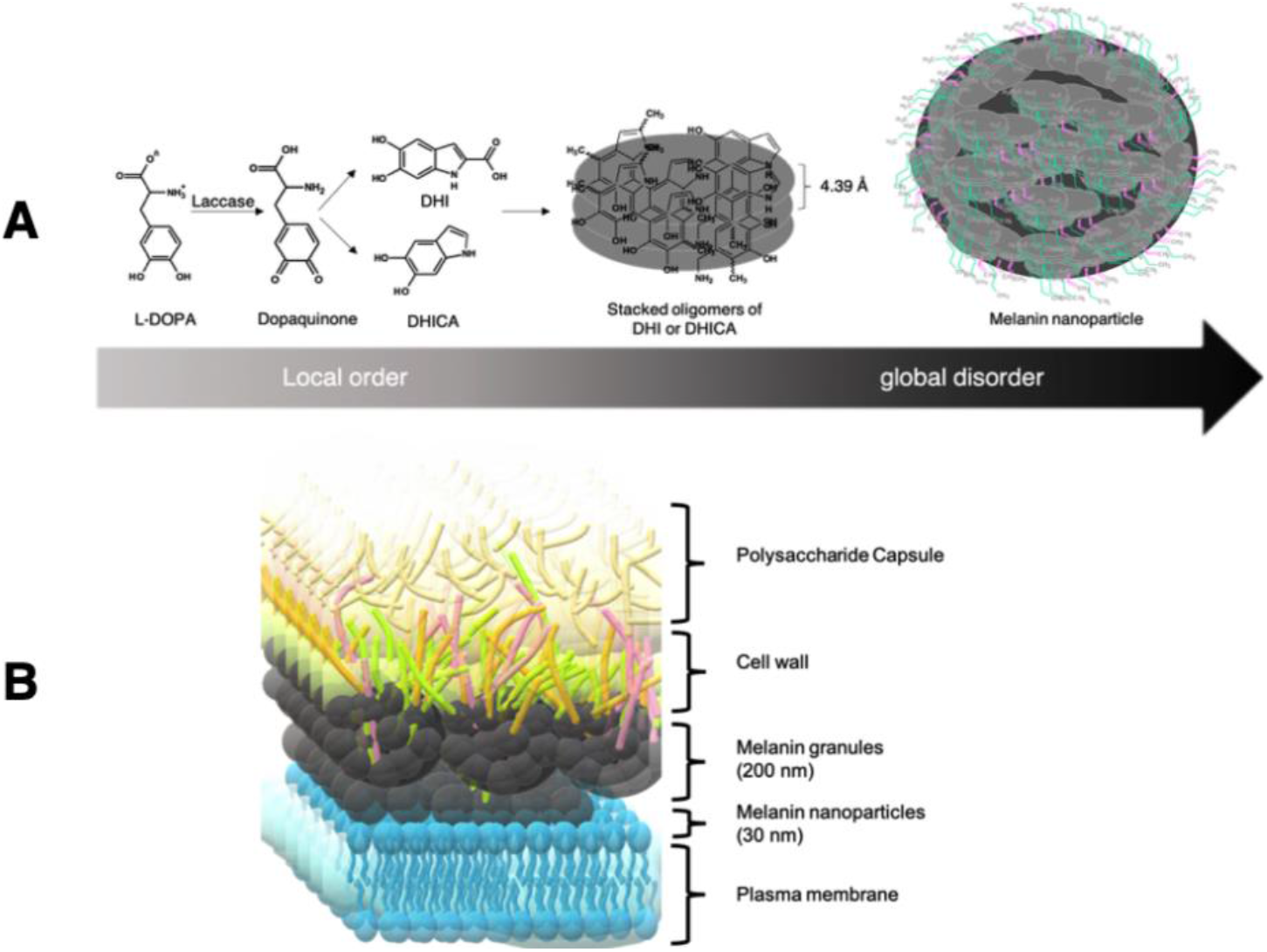
Current views about the synthesis of fungal eumelanins. **A)** Catecholamine precursors like L-DOPA are oxidized by the polyphenol oxidase laccase to form dopaquinone, that undergoes further oxidation to form DHI or DHICA. DHI and DHICA form locally-ordered oligomers that in turn form planar stacks, presumably with stacking distances of about 4.5 Å (44). The higher order of structure of eumelanin is considered ‘disordered’, since different planar structures are oriented in diverse geometries to one another resulting from diversity of DHI/DHICA oligomers stabilized by hydrogen bonding, cation-pi and Van der Waals interactions (48). The aggregation of these simpler units lead to formation of the granular structure that we refer to as melanin granules, which have functional groups that interact with cell wall and cell membrane components such as lipids [(CH_2_)_n_, depicted in green] proteins and polysaccharides (C=C, depicted in pink) (18). **B)** Model representing the architecture of the plasma membrane represented by a lipid bilayer, that interact with melanin nanoparticles that aggregate to form larger melanin granules (21). Melanin granules are anchored to chitin, chitosan and beta-glucan components of the cell wall components that underlie the polysaccharide capsule of *C. neoformans*.

Previous studies from our group have shown that the budding of melanized *C. neoformans* cells involves a decrease in the melanin thickness of the pigmented cell wall near the bud-site (49,50). We have hypothesized that the pigmented nanoparticles released into the extracellular space as part of the normal cell-wall remodeling process are the smallest structural units of melanin found anchored to the cell wall. Hence, we have now investigated the physical properties, supramolecular structure, and protein content of these secreted melanin nanoparticles in the context of current knowledge of the cell-wall associated fungal melanin.

## Results

### Prolonged acid hydrolysis of melanin ghosts yields melanin nanoparticles

Prior studies have shown that the solid yeast-shaped particles lacking all internal structures designated as “melanin ghosts”, are composed of smaller spherical structures ranging in diameter from 40 to 130 nm (21,49). We hypothesized that these particles were held together by non-pigment cellular components, a supposition supported by NMR studies showing that *C. neoformans* ghosts contain tightly associated lipid moieties and cell-wall polysaccharides such as chitin (16,19). Since polysaccharides are typically susceptible to acid hydrolysis, we reasoned that prolonging the hydrochloric acid (HCl) incubation conducted during the melanin ghost isolation protocol would free the smaller spherical melanin particles for further analysis. Consistent with these expectations, increasing the incubation time of the HCl treatment led to significant morphological changes in the melanin ghosts samples. SEM micrographs showed that ghosts, subjected to the prolonged acid incubation, initially appeared to be deformed as the relatively larger melanin particles, located close to the external surface, are release; over time, they took on a “hollowed out” appearance as the smaller melanin nanoparticles were subsequently liberated (**Figures 2A-C**). In addition, the roughness of the ghost surface increased with the prolonged acid treatment, suggesting that the removal of acid-labile non-pigment material exposes more of the underlying melanin particles. SEM visualization of material released during 4 h of HCl boiling revealed melanin aggregates between 60 and 160 nm in diameter size, with the majority measuring around 60 to 80 nm, that were surrounded by smaller granule-like melanin particles with diameters less than 60 nm (**Figures 2D-E**). Cross sectional imaging of this suspended material showed a mix of granular nanoparticles ∼30 nm in diameter with variable electron density and degrees of consolidation (**Figure 2F**). Further TEM analyses of these particles using negative staining, which involves coating the sample with a heavy metal stain that protects its structure from collapsing during drying and thus provides contrast for the detection of structural features, revealed images consistent with the notion that these nanospheres form tightly packed melanin aggregates (**Figure 2G-I**). Altogether, these results established that melanin ghosts of *C. neoformans* are composed of aggregated melanin nanoparticles arranged as small to large sized spherical particles, which are found embedded within an acid-labile matrix that produces a smooth appearance for these cell-sized structures.

**Fig. 2.**
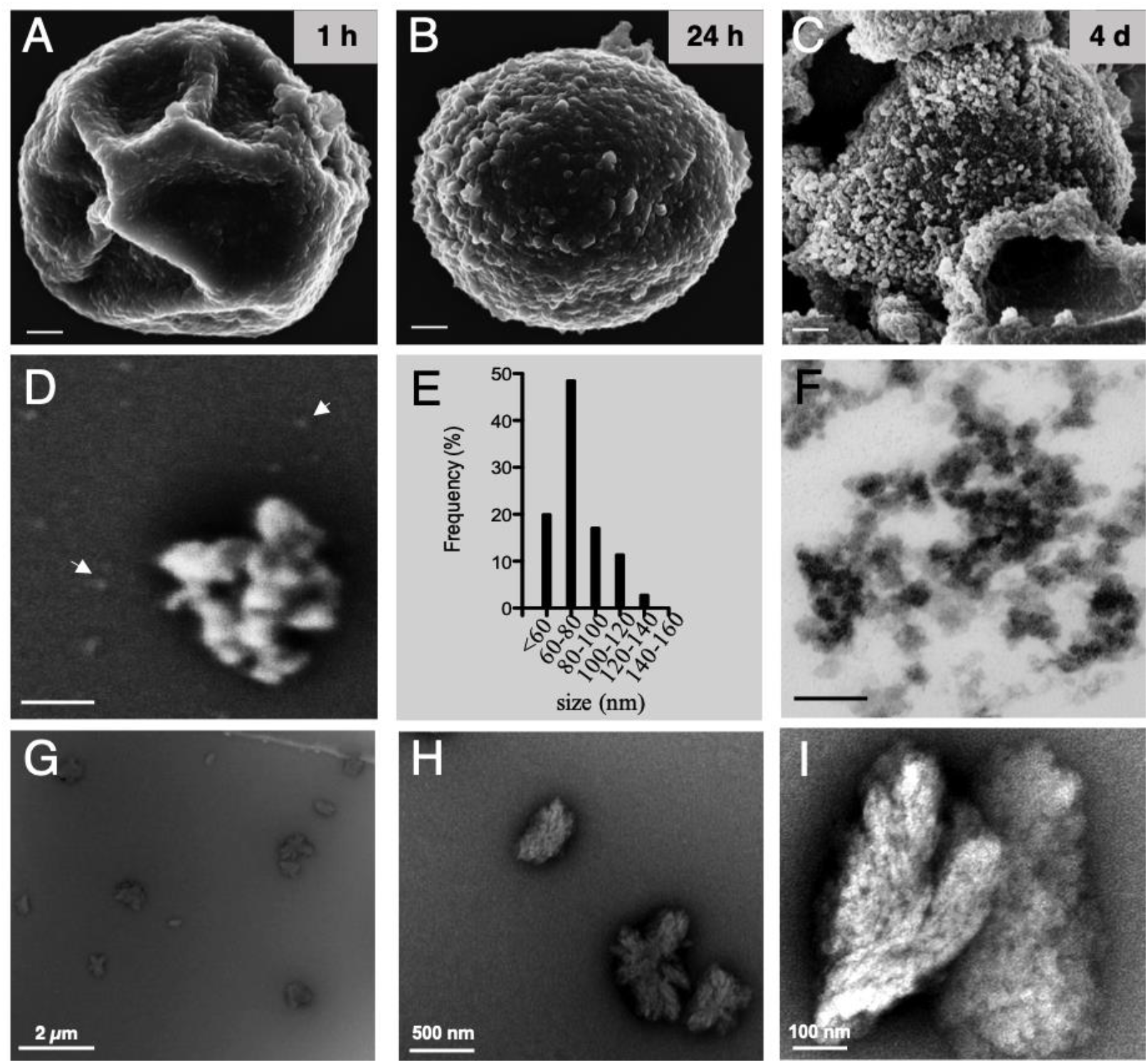
Fungal melanin nanoparticles are released by acid hydrolysis of *C. neoformans* melanin ghosts. SEM images of *C. neoformans* melanized cells subjected to standard melanin isolation but variable incubation time in 6 N HCl for **A)** 1 h (normal), **B)** 24 h, or **C)** 4 days. Particles in solution released during 4 h of acid treatment were analyzed by multiple techniques: **D)** SEM micrographs showing nanoparticles (*arrows*) surrounding them. **E)** Size distribution of released particle population determined by measuring diameters in SEM micrographs. Two hundred particles were counted. **F)** Cross-sectional images of suspended acid-released nanoparticles showing variable electron density and degree of aggregation. Scale bars, 100 nm. **G-I)** Acid-released particles recovered from *C. neoformans* strain 24067 analyzed by TEM using negative staining.

### *C. neoformans*-derived melanin granules differ from auto-polymerized L-DOPA

L-DOPA auto-polymerization occurs both in the presence of *C. neoformans* cells or in cell-free conditions (17). However, in the various cellular systems where melanogenesis has been studied, melanin synthesis occurs within specialized vesicles known as melanosomes (24,27–29,51). During the budding process of melanized fungal yeasts, melanosomes in the cell wall are either degraded and/or laterally displaced so that the daughter cell can emerge (49,50,52). Whereas in synthetic systems, melanin formation is due to the auto-polymerization of precursor molecules, such as L-DOPA. Consequently, we hypothesized that the released melanosomes would be found in the supernatants of *C. neoformans* melanized cultures, possibly in association with extracellular vesicles. The culture supernatants from a *C. neoformans* wild type strain (grown with or without L-DOPA) and a strain defective for melanin synthesis, *ΔLAC1,2* mutant, (**Figures 3A-D,G**) were centrifuged at 100,000 x g and the resulting pellets were analyzed. The solid material from each of the cell cultures, herein refered as crude material, was found to contain particles mostly around 100 nm in diameter as measured by Dynamic Light Scattering (DLS) (**Figures 3B,E,H**). As a control, we also analyzed the pelleted material from a culture containing L-DOPA and no fungal inoculum, observing a population of larger particles from 200 to 300 nm in diameter (**Figure 3K**) that we attributed to melanin aggregates from L-DOPA auto-polymerization. Characterization of these materials by TEM using negative staining was also performed. The sample from *C. neoformans* grown with L-DOPA revealed the presence of electron-dense aggregated granular structures, henceforth called melanin granules (**Figure 3C**). Similar aggregated structures were seen in the Δ*LAC1,2* strain sample (**Figure 3I**), while they were absent in the wild type without L-DOPA (**Figure 3F**). In the L-DOPA sample without *C. neoformans* cells we observed irregular and non-spherical aggregates that did not resemble the melanin granules observed for the *C. neoformans* cultures (**Figure 3L**), and which we attribute to L-DOPA auto-polymerization.

**Fig. 3.**
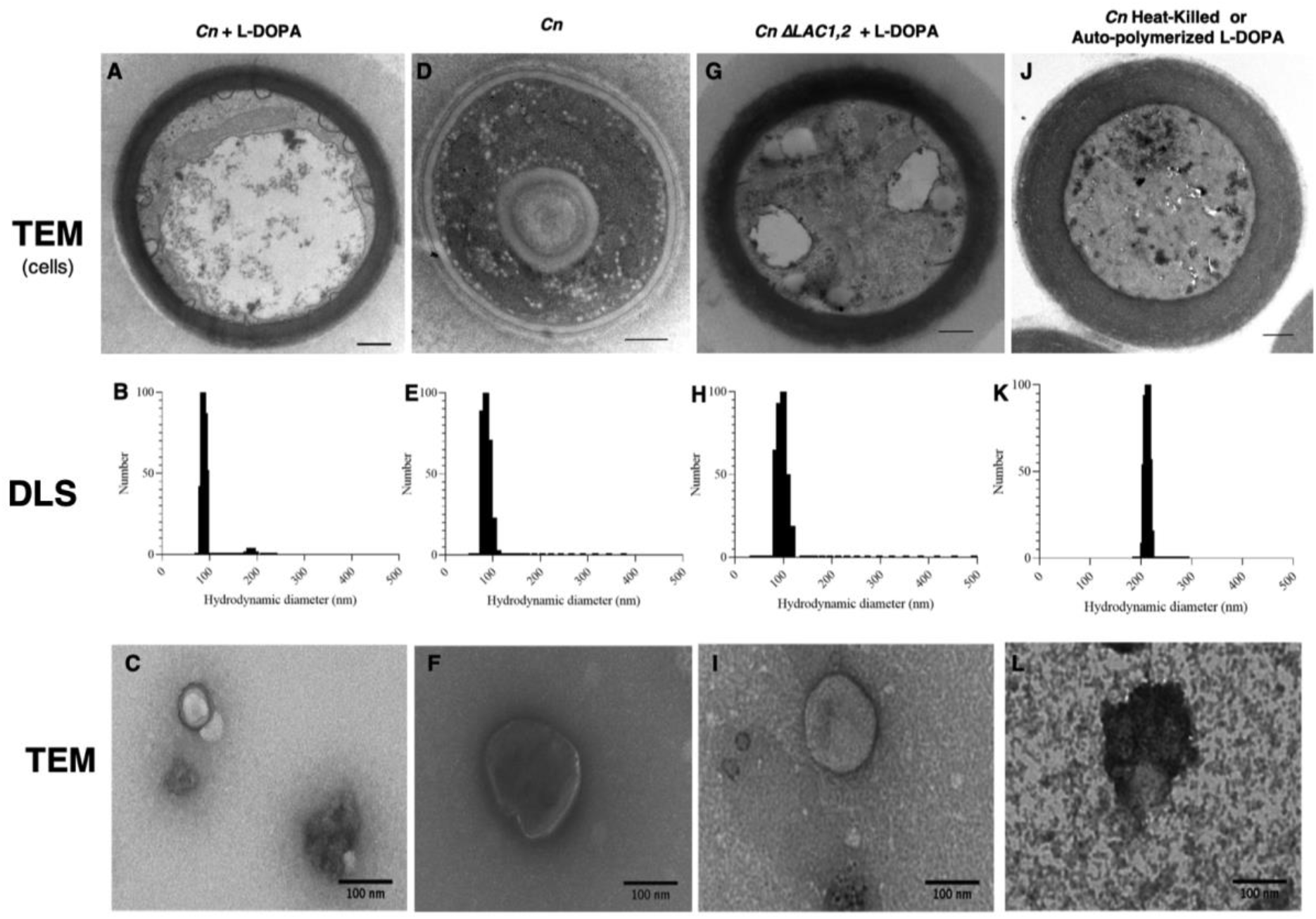
Secreted vesicles and melanin granules from *C. neoformans* are found in culture supernatants. *Top panel*, Cross sectional view of representative cells by TEM. Scale bars, 500 nm. *Middle panel*, Hydrodynamic diameter of vesicles and melanin granules measured by DLS. *Bottom panel*, Representative micrographs by TEM of vesicles and melanin granules (often present as aggregates) using negative staining **A-C)** *C. neoformans* grown in MM with L-DOPA. **D-F)** *C. neoformans* grown in MM without L-DOPA. **G-I)** *C. neoformans* Δ*LAC1,2* strain grown in MM with L-DOPA. **J)** *C. neoformans* heat-killed grown MM with L-DOPA. **K-L)** MM with L-DOPA. Data represent from three independent experiments.

Melanin has the unique ability to absorb almost every wavelength of light and in contrast to most other natural chromophores, has a broadband monotonic absorption spectrum (53,54). Thus, we used UV-Vis absorbance spectroscopy to verify whether the aggregated granular particles visualized by TEM are indeed melanin pigments. A monotonic curve was observed for the pelleted cellular material collected from *C. neoformans* cells grown in MM with L-DOPA (melanin granules and vesicles), but not for Δ*LAC1,2* strain grown in MM with L-DOPA (L-DOPA aggregates and vesicles) nor for the fungal cells grown in MM lacking L-DOPA (vesicles) (**Figure 4A**). DLS also yields a dispersity index, which is a measure of the dispersion (or spread) in the estimated hydrodynamic sizes of colloidal particles in a solution (15, 16). We observed that melanin granules had lower dispersity than auto-polymerized L-DOPA (**Figure 4B**). Furthermore, negative staining TEM allowed us to visualize spherical nanoparticles ∼200 nm in diameter, which were isolated from the supernatant of *C. neoformans* melanized cultures (**Figures 5A,B**) whereas, melanin resulting from auto-polymerized L-DOPA particles, showed minuscule structures with no specific pattern for their association (**Figures 5C,D**). These results demonstrated that *C. neoformans* extracellular vesicles along with melanin granules are found in the culture supernatant, while aggregated material seen in the Δ*LAC1,2* strain have similar aggregation properties to auto-polymerized L-DOPA. Thus, melanin granules have unique colloidal characteristics; and these particles are distinct structures in size and morphology distinguishable from melanin resulting from auto-polymerized L-DOPA

**Fig. 4.**
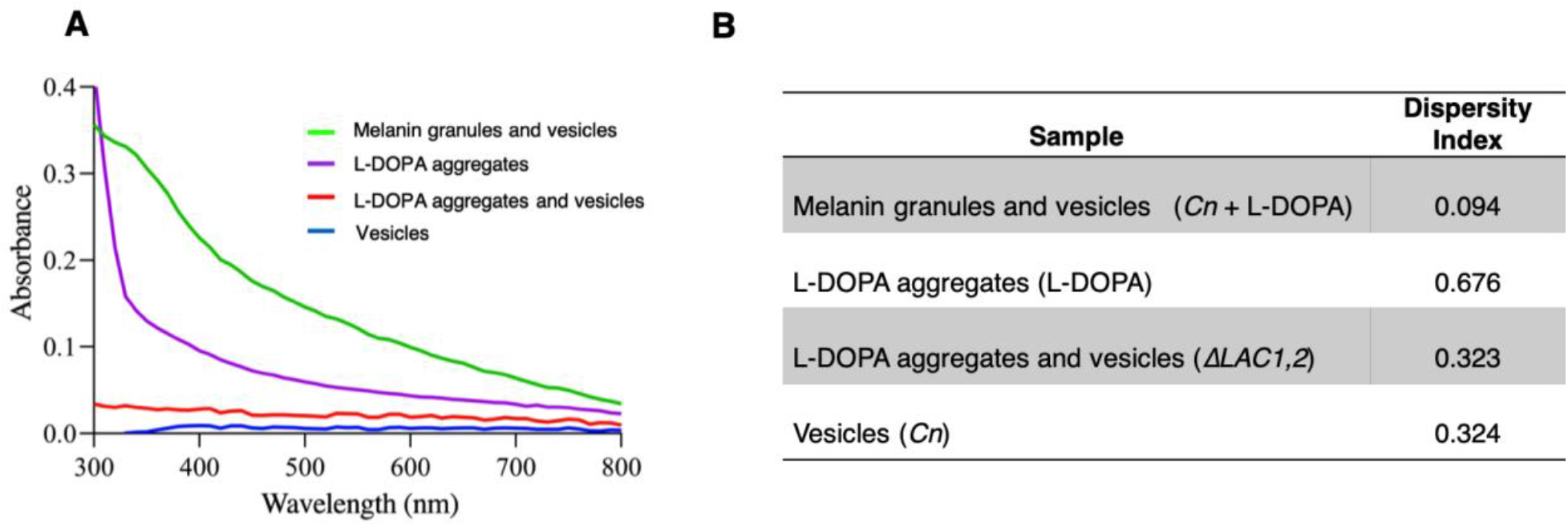
Secreted vesicles and melanin granules from *C. neoformans* are biologically synthesized. **A)** Absorbance spectra of vesicles and melanin granules. Melanin granules and vesicles collected from *C. neoformans* grown in MM with L-DOPA (orange) has a broadband optical absorption curve characteristic of melanin. A curve for auto-polymerized L-DOPA is apparent in MM (purple), and no such curve is visible for vesicles collected from *C. neoformans* in MM lacking L-DOPA or for material collected from *C. neoformans* Δ*LAC1,2* in MM with L-DOPA. **B)** Dispersity index values of vesicles and melanin granules. Melanin granules in *C. neoformans* + L-DOPA showed low dispersity, suggesting that the population of melanin granules is highly monodisperse. L-DOPA aggregates had a broad polydisperse distribution. Representative data from two independent experiments.

**Fig. 5.**
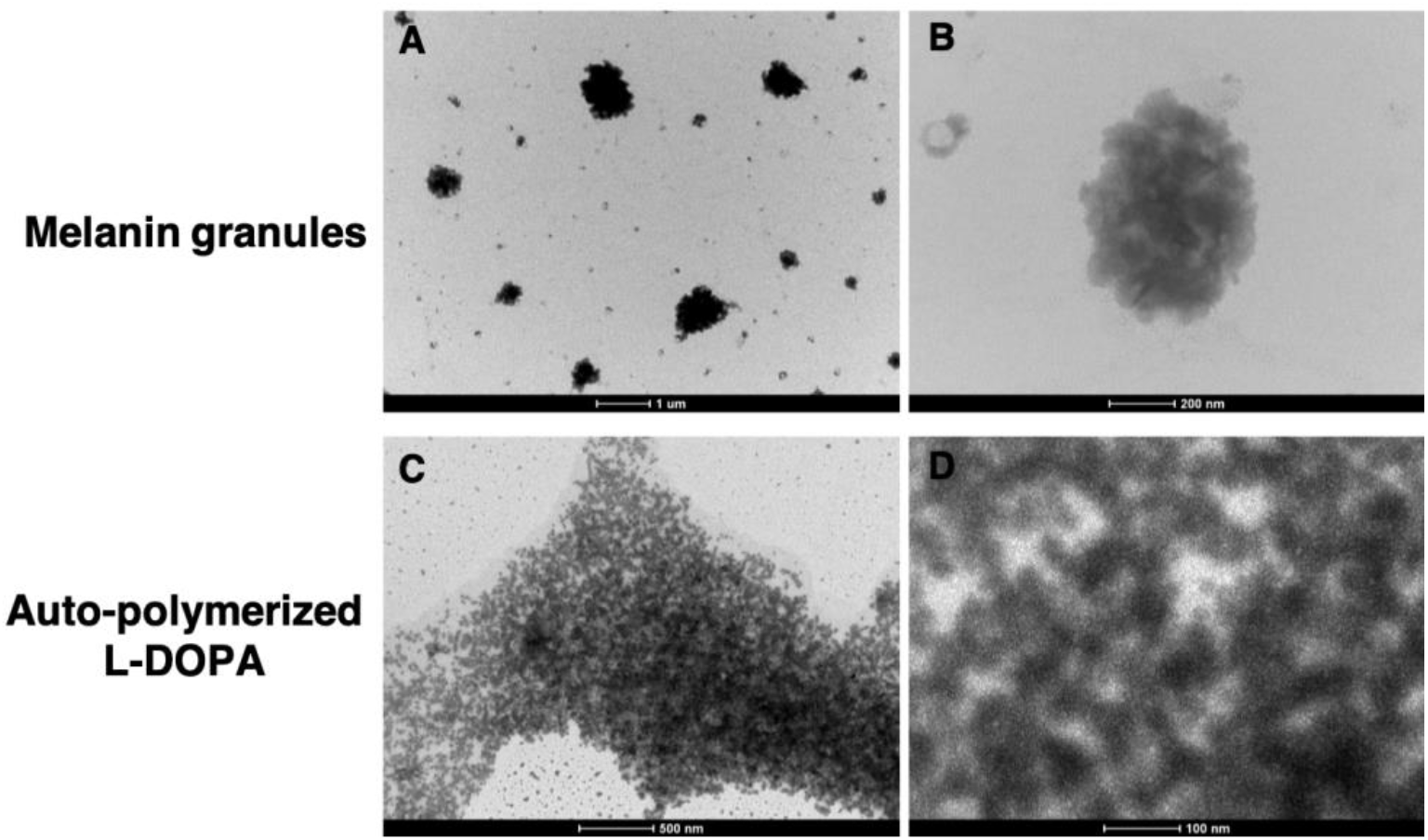
*C. neoformans* melanin granules are shown as highly monodisperse spherical nanoparticles. High resolution TEM micrographs using a 1% phosphotungstic acid (PTA). **A,B)** Melanin granules found along with extracellular vesicles display a spherical shape. **C,D)** Broadly polydisperse minute spherical L-DOPA particles resulted from auto-polymerization. Representative images of three experiments.

### Divalent cations promote aggregation of *C. neoformans* melanin granules

We studied the hydrodynamic size of melanin granules suspensions in the presence of mono (Na^+^) and divalent (Ca^2+^) salt concentrations to gain insight into their intramolecular interactions. The crude material recovered from the culture supernatants of *C. neoformans* cells grown with L-DOPA containing both melanin granules and vesicles (hereinafter referred to as crude melanin granules) aggregated at >0.01 M CaCl_2_, >0.1 M NaCl, and >10X PBS (**Figure 6A-C**). These results indicate that divalent cations promote melanin aggregation; or fusion events at lower concentrations than monovalent cations. Alternatively, divalent cations likely promote particle aggregation by neutralizing negative charges on the surface of melanin (55).

**Fig. 6.**
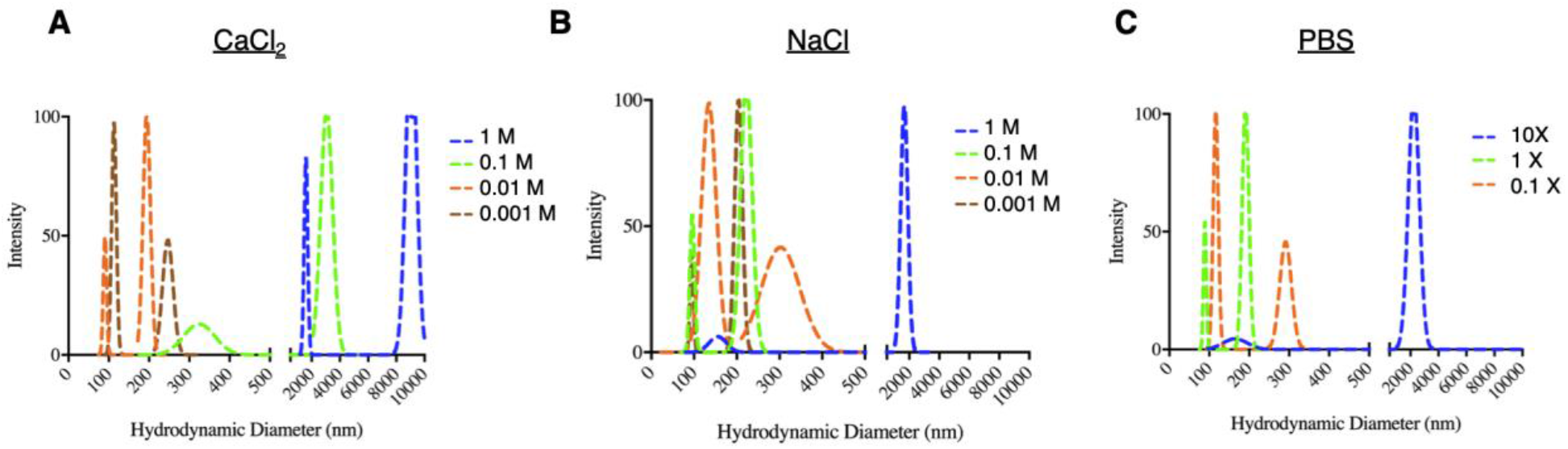
Divalent cations caused the aggregation of crude melanin granules at lower concentrations than monovalent cations. Size distribution of melanin granules and vesicles collected from *C. neoformans* grown in MM with L-DOPA resuspended in **A)** 1 M, 0.1 M, 0.01 M, 0.001 M CaCl_2_; **B)** 1 M, 0.1 M, 0.01 M, 0.001 M NaCl; and **C)** 10X, 1X, 0.1X PBS. Representative data from two independent experiments.

### *C. neoformans* melanin granules can be isolated based on their buoyancy

Ultracentrifugation of *C. neoformans* cell culture supernatants leads to the recovery of a mixture of vesicles and protein aggregates (56). We hypothesized that in addition to these constituents, the cellular material recovered from melanized cell cultures would contain tightly packed melanin granules of high molecular weight and low volume that could be separated from the vesicles and protein aggregates by exploiting their buoyancy. Thus, we analyzed the pelleted crude material from the supernatants of *C. neoformans* non-melanized and melanized cultures using density gradient ultracentrifugation. A distinct dark band visible by examination with the naked eye was apparent towards the bottom of the tube for the melanized sample, corresponding to fraction 5 (**Figure 7A**). TEM imaging of the collected fractions revealed vesicles primarily in fractions 1 and 2, for both samples (**Figure 7B**); aggregated electron dense nanoparticles ∼30 nm in diameter, corresponding to the dimensions and appearance of melanin granules, were also apparent in fraction 5 of the melanized sample (**Figure 7B**).

**Fig. 7.**
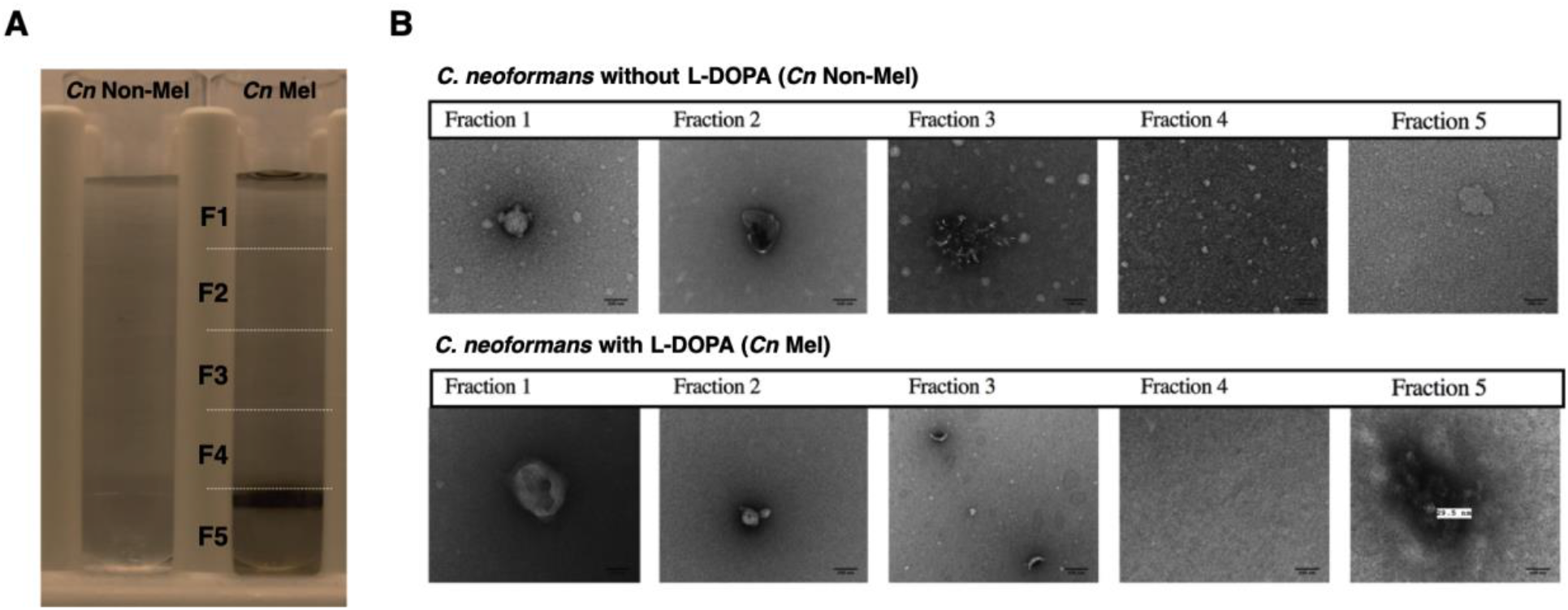
Density gradient separation of melanin granules from extracellular vesicles. **A)** Image of OptiPrep™ density gradient of ultrapelleted culture supernatants (crude material) from *C. neoformans* non-melanized and melanized cultures grown in MM without or with L-DOPA, respectively. **B)** TEM images depicting Fractions 1-5 collected from density gradient centrifugation. Melanin granules are visible in fraction 5 of the *C. neoformans* melanized sample. Representative data from three independent experiments. Scale bars, 100 nm.

### Melanin particles from ghosts and culture supernatants are similar

We entertained the hypothesis that melanin granules in *C. neoformans* culture supernatants were similar to those recovered from prolonged acid hydrolysis of melanin ghosts. Consequently, we sought to mechanically break down the melanin ghosts isolated from *C. neoformans* by extended ultrasonication, avoiding the chemical exposure. We found that ultrasonication of melanin ghosts for up to 12 min resulted in the release of particles in the range 0-200 nm measured by DLS (**Figure 8A**). SEM visualization revealed that an ultrasonic pulse as short as a 30 s led to fracture of the melanin ghosts characterized by jagged edges, which then turned into smaller round structures exhibiting a coarse and rough appearance as the process continued (**Figure 8B**). The spherical melanin particles that resulted from extensive physical disruption of *C. neoformans* melanin ghosts showed a similar size distribution to the crude melanin granules in the culture supernatant. These data support the proposal that *C. neoformans* cell-wall melanin is composed of tightly packed melanin granules with a common substructure of melanin nanoparticles, which are also release to the extracellular environment.

**Fig. 8.**
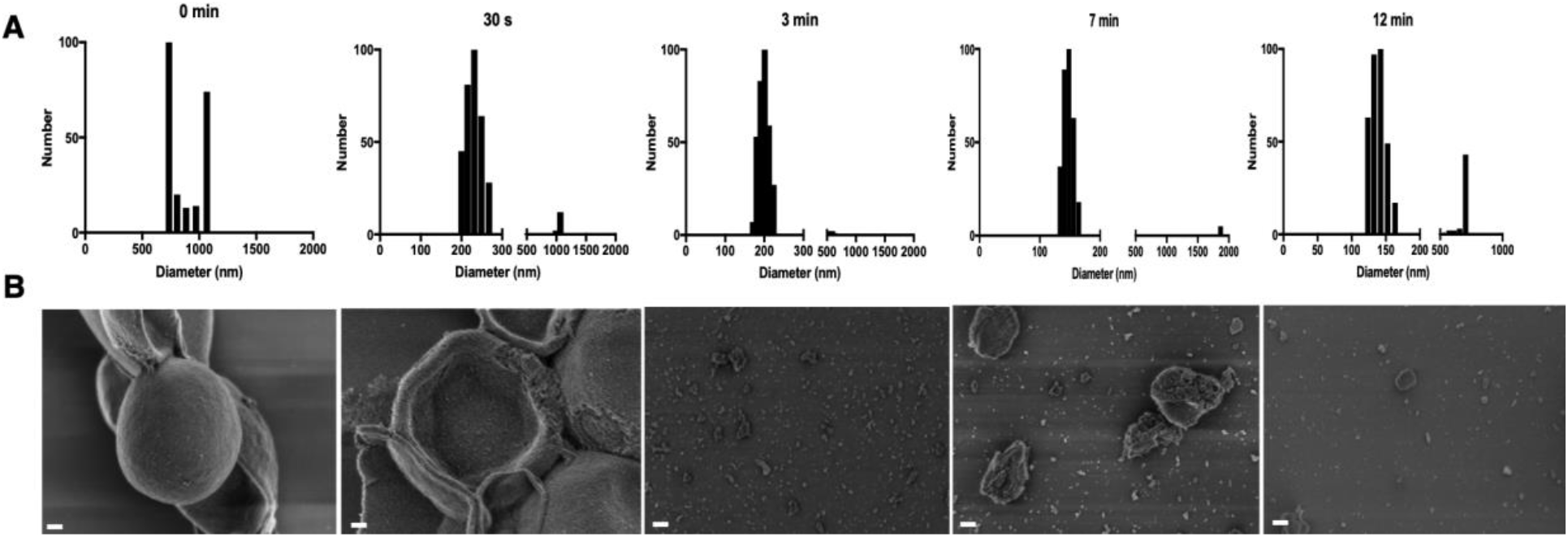
Sonic cavitation of melanin ghosts from *C. neoformans* reduces melanin to particles of ∼ 200 nm. **A)** Size distribution of melanin particles by dynamic light scattering as sonication time increases. **B)** SEM micrographs showing melanin break down. Scale bar, 200 nm. Representative micrographs of 3 independent experiments.

To gain further insight into the structure of *C. neoforman*s melanin granules we used cryoEM. Micrographs of fungal melanin granules isolated by buoyancy revealed a pattern of similarly sized, round beads on a string (**Figure 9A**). Each spherical nanoparticle, ∼200 nm in diameter, resembles a multilayered structure with well-defined edges but not distinguishable lipid bilayer, suggesting that these structures may result from stepwise aggregation of smaller particles (**Figure 9B**). We considered whether the structures visualized by cryo-EM images were water ice particles. We cannot unequivocally distinguish between these possibilities by this form of microscopy. Nevertheless, we noted that our putative melanin structures are relatively uniform, have dimensions comparable to those obtained melanin by other EM methods (**Figure S1**), are translucent, less electron dense than ice, and do not express the geometric structures associated with water ice. Hence, we are confident that these are melanin granules.

**Fig. 9.**
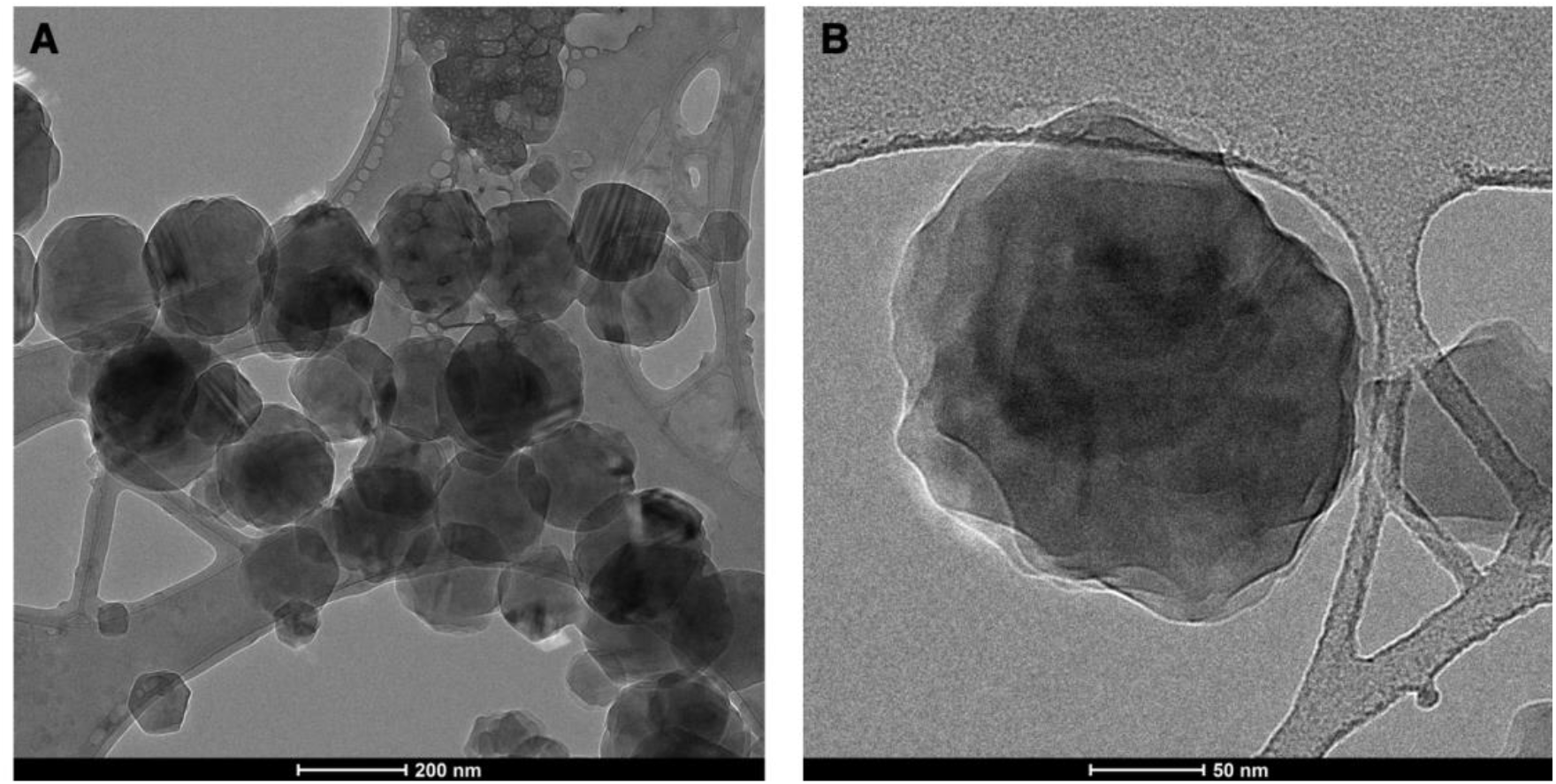
Ultrastructure of *C. neoformans* melanin granule suggests a multilayered superstructure. **A)** CryoEM micrographs exhibit melanin granules held together as beads on a string. **B)** Spherical structure representative of the fundamental unit of *C. neoformans* cell-wall melanin ∼200 nm in diameter.

### Melanin granules from *C. neoformans* exhibit a characteristic EPR signal for eumelanins

A key feature of melanins is the presence of a stable free-radical population that can be detected by EPR (57), which serves as a molecular fingerprint for natural materials such as charcoal (58). Eumelanins can be identified based on the modification of their EPR signals by a number of agents including light, pH, redox agents, and diamagnetic multivalent metal ions (40). The crude melanin granules recovered from the *C. neoformans* culture supernatant yielded a single slightly asymmetric EPR trace similar to the spectrum previously described for melanin ghosts (59) (**Figure 10A**). Irradiation of fungal crude melanin granules with a 250 W LED white light enhanced the EPR signal intensity 1.2 times, while doping with zinc ions increased it 5.3-fold. Alkaline pH had little or no effect on the EPR signal of crude melanin granules, but low pH was found to attenuate their EPR signal intensity, possibly due to changes in protonation of accessible ionizable groups. The melanin ghosts EPR signal was insensitive to pH (**Figure 10B**). Thus, these results demonstrated that melanin granules found in the culture media of *C. neoformans*, in presence of L-DOPA, retain the paramagnetic properties characteristic of eumelanins.

**Fig. 10.**
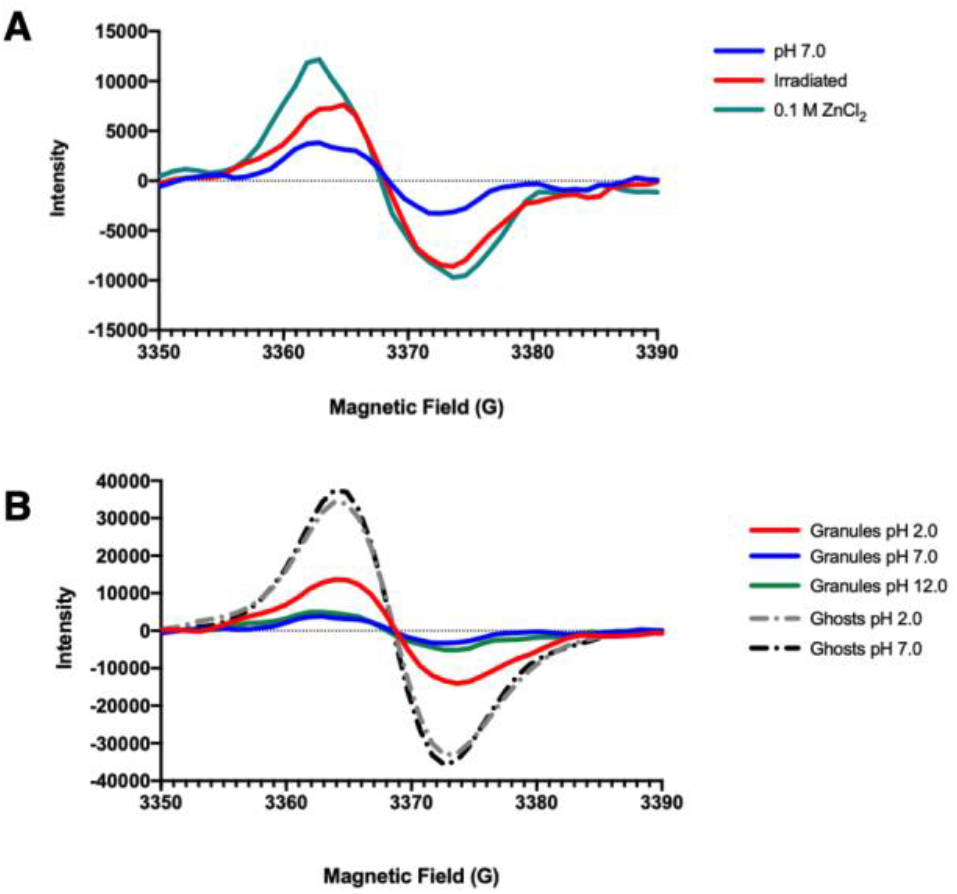
Electron paramagnetic resonance spectroscopy analysis of *C. neoformans* crude melanin granules. **A)** EPR of *C. neoformans* crude melanin granules suspended in distilled water at pH 7 (blue); after irradiation for 20 min with white light from a 250-W LED white lamp; and suspended in a solution of 0.1 M ZnCl_2_. **B)** EPR of *C. neoformans* crude melanin granules suspended in distilled water at acidic pH adjusted with 1 N HCl (red); suspended in distilled water at pH 7 (blue); suspended in distilled water at basic pH adjusted with 1N NaOH (green).

### *C. neoformans* melanin is molecularly associated with non-polysaccharide cellular constituents

Previous ssNMR studies by our group have revealed that melanin ghosts isolated from *C. neoformans* cells are an assembly of the L-DOPA-derived melanin pigment and the cellular “scaffolding” moieties on which it the pigment is deposited (16). Here we again used ssNMR to determine whether the melanin granules that are present in the pelleted crude material from *C. neoformans* culture supernatants are also associated with non-pigment cellular constituents. To obtain a sample mass sufficient for ssNMR analysis, we took advantage of the previously reported *C. neoformans* mutant strain ST211A (GenBank Accession Number MK609896), which displays a “leaky melanin” phenotype; “leaky melanin” strain cells are able to synthesize melanin, but the cell wall is unable to the retain the pigments and thus they leak into the extracellular space (33). The ^13^C cross-polarization magic angle spinning (CPMAS) spectrum of the crude melanin granules obtained from the supernatant of a *C. neoformans* ST211A cell culture containing uniformly ^13^C-enriched glucose as the sole carbon source and L-DOPA (at natural abundance) is compared to that of isolated melanin ghosts in **Figure 11 (A,B)**; this isotopic labeling scheme allows us to selectively observe melanin-associated cellular components without interfering signals from the melanin pigment. Whereas the most prominent peaks in the melanin ghosts CPMAS spectrum are centered at ∼30 ppm, likely arising from the aliphatic “tails” of vesicle-derived lipids functionalized during melanization, the crude melanin granules spectrum is dominated by signals in the region between ∼55 and 105 ppm that correspond to various types of polysaccharide ring carbons. The strong presence of polysaccharides in this crude preparation is unsurprising since our research group has previously demonstrated that capsular polysaccharides are synthesized intracellularly and transported to the extracellular space within vesicles (26). To test whether these polysaccharides are indeed capsular components that are not associated with the melanin granules, the crude sample was reexamined after 30 minutes of incubation in hot HCl (**Figure 11C**). This relatively short treatment was sufficient to drastically diminish the polysaccharide content so that the signals in the ∼55-105 ppm region were of lesser relatively intensity than observed for melanin ghosts, which had been subjected to an hour of HCl boiling in addition to other enzymatic and chemical treatments during sample preparation.

**Fig. 11.**
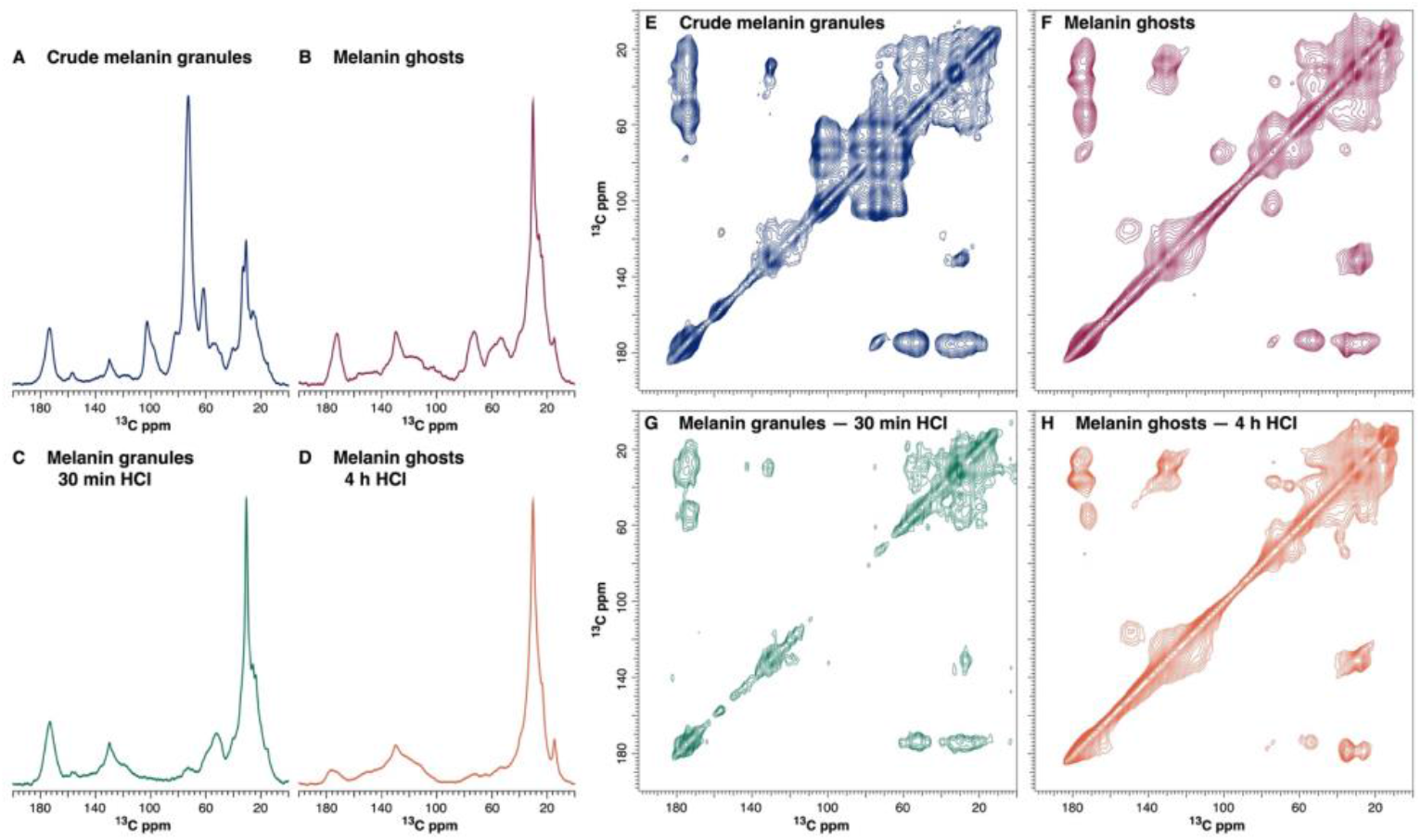
ssNMR analyses of *C. neoformans* crude melanin granules demonstrated a weak association to polysaccharides and suggested the presence of peptides or proteins strongly “functionalized” during melanization. **A,B)** ^13^C cross-polarization magic-angle spinning (CPMAS) spectra of crude melanin granules and melanin ghosts isolated from the leaky-melanin strain *C. neoformans* ST211A, showing dominant signals for polysaccharides (∼55-105 ppm) or lipids (∼30 ppm), respectively. **C,D)** ^13^C CPMAS spectra of crude melanin granules and melanin ghosts post-acid treatment revealed that lipid signal (∼30 ppm) corresponding to fatty acids is intimately associated with melanin particles. **E,F)** 2D ^13^C-^13^C DARR spectra of crude melanin granules confirmed augmented polysaccharide content (region ∼55-105 ppm) whereas in melanin ghosts these constituents are less prominent. **G,H)** 2D ^13^C-^13^C DARR spectra of crude melanin granules and melanin ghosts post-acid treatment revealed that both samples are composed of the same cellular constituents and suggest the presence of lipids and possibly protein remnants (∼53×175 ppm) that might be strongly associated with the melanin polymer.

Further insight into the molecular identity of the cellular constituents that associate with melanin granules was obtained from 2D ^13^C-^13^C DARR spectra that display correlations between proximal carbon pairs equivalent to approximately 1-2 bond-lengths in distance. In analogy to our ^13^C CPMAS data, the DARR plot of the crude melanin granules displays a multitude of prominent cross-peaks corresponding to correlations between spatially-close polysaccharide ring carbons (∼55-105 ppm), whereas this region in the melanin ghosts 2D plot only contains a handful of correlations that we have previously attributed to the cell-wall scaffolding moieties (e.g., chitin) on which the pigment is deposited (15) (**Figure 11E,F**). No cross-peaks attributable to polysaccharides were observed for the crude melanin granules after the 30-minute incubation in hot HCl, supporting our hypothesis that the polysaccharides were indeed superficially bound capsular components and that melanin granules released into the extracellular space are not tightly associated with cell-wall constituents as for melanin ghosts (**Figure 11G**). Moreover, all of the spectral features displayed in the 2D DARR plot of the HCl-treated melanin granules were also observed in the melanin ghosts spectrum. We interpreted this spectroscopic similarity to indicate that the 30-minute HCl incubation was adequate to hydrolyze all cellular moieties that were not intimately associated with the melanin granules. The major difference between the two datasets is the presence of cross-peaks attributable to the polysaccharide scaffolding constituents in the melanin ghost DARR spectrum. To test the robustness of the association between these residual polysaccharides and the melanin pigment, we subjected the melanin ghosts to a series of treatments in boiling HCl using incrementally longer incubation times and monitored the peak intensities of the non-pigment constituents using ^13^C CPMAS experiments (**Figure S2**). Increasing the incubation time from 2 h to 4 h resulted in only minimal spectral differences, and thus we chose to further examine the 4 h HCl-treated melanin ghosts by acquiring a 2D DARR spectrum (**Figure 11H**). Although the additional 4 h HCl treatment was effective in diminishing the signals corresponding to polysaccharides to undetectable levels (**Figures 11 F,H**), the majority of other cross-peaks were preserved; furthermore, these cross-peaks were also retained in the DARR spectrum of the crude melanin granules after 30 min of HCl treatment (**Figure 11G**). This comparison indicates that the two samples contain many of the same cellular constituents, which are likely strongly associated with the pigment and could play a role in melanin ultrastructure. Interestingly, the correlations common to both sample spectra suggest the presence of lipids and likely also protein remnants: the cluster of signals centered around ∼53 x 175 ppm is consistent with correlations between the alpha- and carbonyl-carbons of amino acids within a peptide bond. However, this later assignment is provisional and requires further testing.

### Melanin-associated proteins are identified from *C. neoformans* melanin granules

To assess the possibility that *C. neoformans* melanin nanoparticles are associated with, or possibly covalently bonded to, melanization-specific proteins, we performed a proteomic analysis on the isolated melanin granules comprising the visibly dark band (fraction 5) after density gradient centrifugation. To distinguish between proteins that are vesicle-associated and/or loosely bound from those chemically functionalized by the melanin nanoparticles, we included as controls fraction 5 samples collected from a *C. neoformans* culture grown in MM without L-DOPA as well as auto-polymerized L-DOPA. To ensure that sufficient material was available for LC-MS/MS analysis, 2-3 density gradients procedures were performed for each sample and their respective fractions 5 were pooled together. Melanin granule-associated proteins were solubilized, reduced, carboxymethylated, and digested before LC-MS/MS measurements. Data analysis yielded was the consistent identification of four proteins exclusively associated with *C. neoformans* melanin granules (**Table 1**). A bioinformatics analysis to examine the presence of predicted signal peptides and GPI anchors showed that three proteins had an N-terminal signal peptide, two of which (Cig1 and Blp1), were also found in extracellular vesicles (60). Only one protein, CNAG_05312, had both an N-terminal signal peptide and a GPI-anchor indicative of plasma membrane association. These secreted peptides were previously reported to play critical roles in *C. neoformans* virulence (61–65). Taken together, these results revealed that *C. neoformans* melanin granules are indeed intimately associated with proteins that employ both a conventional and non-conventional secretory pathway for regulation, and which may be involved with structural and/or functional properties of the cryptococcal eumelanin.

**Table 1.**
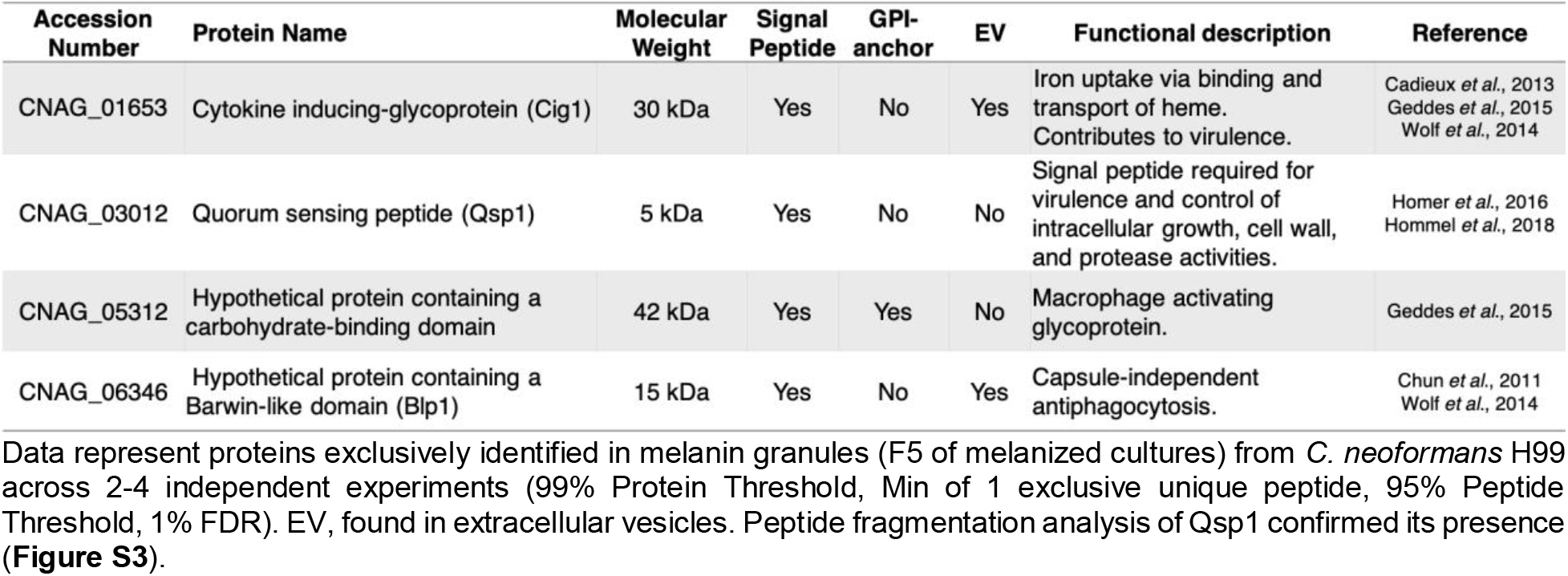
Identified proteins associated to *C. neoformans* melanin granules

| Accession Number | Protein Name | Molecular Weight | Signal Peptide | GPI-anchor | EV | Functional description | Reference |
| --- | --- | --- | --- | --- | --- | --- | --- |
| CNAG_01653 | Cytokine inducing-glycoprotein (Cig1) | 30 kDa | Yes | No | Yes | Iron uptake via binding and transport of heme. Contributes to virulence. | Cadieux *et al.*, 2013<br>Geddes *et al.*, 2015<br>Wolf *et al.*, 2014 |
| CNAG_03012 | Quorum sensing peptide (Qsp1) | 5 kDa | Yes | No | No | Signal peptide required for virulence and control of intracellular growth, cell wall, and protease activities. | Homer *et al.*, 2016<br>Hommel *et al.*, 2018 |
| CNAG_05312 | Hypothetical protein containing a carbohydrate-binding domain | 42 kDa | Yes | Yes | No | Macrophage activating glycoprotein. | Geddes *et al.*, 2015 |
| CNAG_06346 | Hypothetical protein containing a Barwin-like domain (Blp1) | 15 kDa | Yes | No | Yes | Capsule-independent antiphagocytosis. | Chun *et al.*, 2011<br>Wolf *et al.*, 2014 |

Based on the results obtained in this study, we propose a model of *C. neoformans* cell wall melanization that begins intracellularly and requires melanosome transport to the cell wall for their consolidation into melanin granules (**Figure 12**). Melanin synthesis occurs within melanosomes (nanospheres 10-20 nm in diameter) enclosed in multivesicular bodies (MVBs) (∼450 nm in diameter) (**Figure 12A**) and laccase-containing vesicles (∼80 nm in diameter) (**Figure 12B, upper inset**). These organelles may be signaled by the Qsp1 mature peptide to activate melanogenesis (64). Ongoing melanization is evidenced by the presence of melanosomes with variable degrees of aggregation and electron density within these double-layered compartments. The melanosomes aggregate into ∼60 nm particles (**Figure 12B, lower inset**), both within MVBs and in the cell cytoplasm, and are eventually transferred into the cell wall by events that are facilitated by the presence of invaginations in the plasma membrane (**Figures 12C,D**). These structures are characterized by finger-like protrusions that “scoop” melanosomes from the cytosol to transfer them to the cell wall. Further aggregation of the melanosomes into melanin granules (up to 200 nm in diameter) and anchoring within the cell wall could be augmented by vesicle materials, such as laccase and Blp1, respectively. Some of these vesicles are delivered to the cell wall within MVBs that fuse with the plasmatic membrane (**Figures 12E**). Melanin granules’ concentric arrangement within the cell wall uses as a scaffold membranous sheets shown as multilaminated structures of cryptococcal cell walls (**Figures 12F**). Finally, fungal melanosomes are held together by and anchored to cell-wall constituents (i.e. chitin, chitosan, glucans) due to diverse interactions: i) associative or covalent interactions between melanin-associated proteins and cell-wall components, and ii) electrostatic interactions of oppositely charged melanin-chitosan pairs. As the fungal cell replicates, cell wall remodeling is required to allow budding; thus the melanized scaffold is locally modified by secreted enzymes and allows the release of melanin granules (**Figure 12G**) into the surrounding environment. A summary of *C. neoformans* melanin particles analyzed with multiple microscopic techniques is shown in **Figure 13**.

**Fig. 12.**
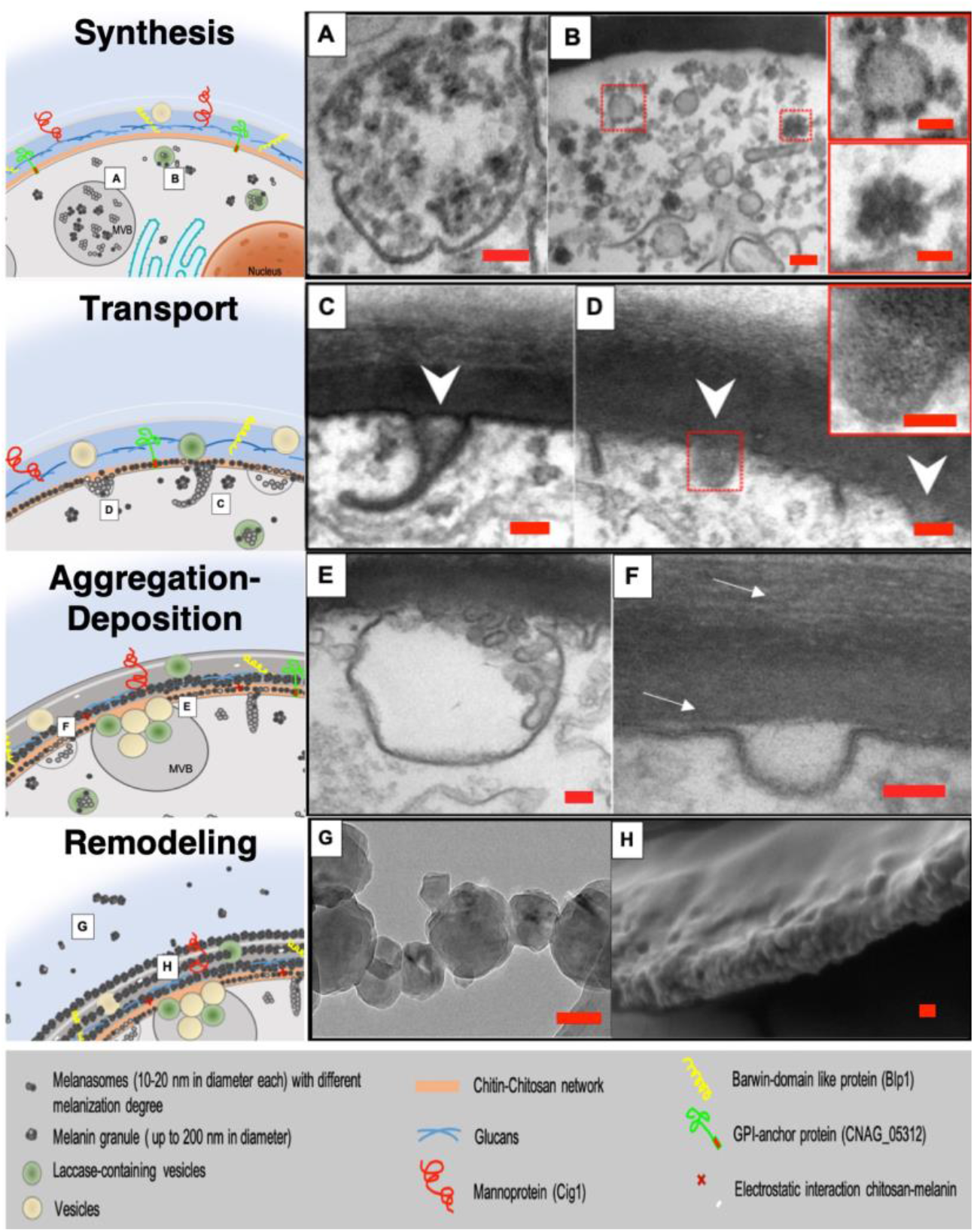
Illustrative model of melanization in *C. neoformans*. Melanogenesis in *C. neoformans* is characterized by multiple events that can be grouped into four phases: synthesis, transport, aggregation-deposition, and remodeling. Intracellular biosynthesis of the polymer occurs within melanosomes (nanospheres 10-20 nm in diameter) enclosed in multivesicular bodies (MVBs) (60,81) (**A**) and laccase-containing vesicles (89) (**B, upper inset**). These organelles could be signaled to start the biopolymer production via incorporation of the mature Qsp1 peptide (64). Active melanization is evidenced by the presence of melanosomes with variable degrees of pigmentation within these lipid bilayer compartments. Melanosomes forming aggregates ∼60 nm in diameter are also noticeable in the cell cytoplasm (**B, lower inset**). Melanosomes are transported to the cell wall using an unconventional mechanism that involves invaginations of the plasma membrane (79,80,90). Finger-like protrusions of the plasma membrane scoop up melanosomes to transport them from the cytosol to the cell wall (**C,D, white arrow heads**). Once in the cell wall, aggregation of melanosomes into up to 200 nm melanin granules might be promoted by laccase enzyme delivered to the cell wall via MVB fusion with the plasma membrane (46,81,89) (**E**), while their arrangement within the fungal cell wall may use as a scaffold concentric membranous sheets reported in basidiomycetes (49,93,94) (**F, white arrows**). Melanosomes anchor to the cell wall via multiple interactions: i) associative and covalent interactions between them and cell-wall constituents (chitin, chitosan, glucans, lipids), possibly mediated by Blp1-chitin and CNAG_05312-glucans/chitin/chitosan (16,18,19), and ii) electrostatic interactions modulated by chitosan-melanin charges (14,31,32,34). Regular cell growth requires the cell-wall remodeling to allow budding inducing the melanin scaffold degradation/alteration via secretion of of peptidases (96), chitinases (97), and glucanase (62) thus releasing melanin granules and strung-like granules (**G**) into the extracellular environment. Removal of non-pigmented acid-labile cell-wall components exposes the concentric layered arrangement of the melanin granules within the cryptococcal cell wall from melanin ghosts (48,94) (**H**). Scale bars, 100 nm (main micrographs), and 50 nm (insets).

**Fig. 13.**
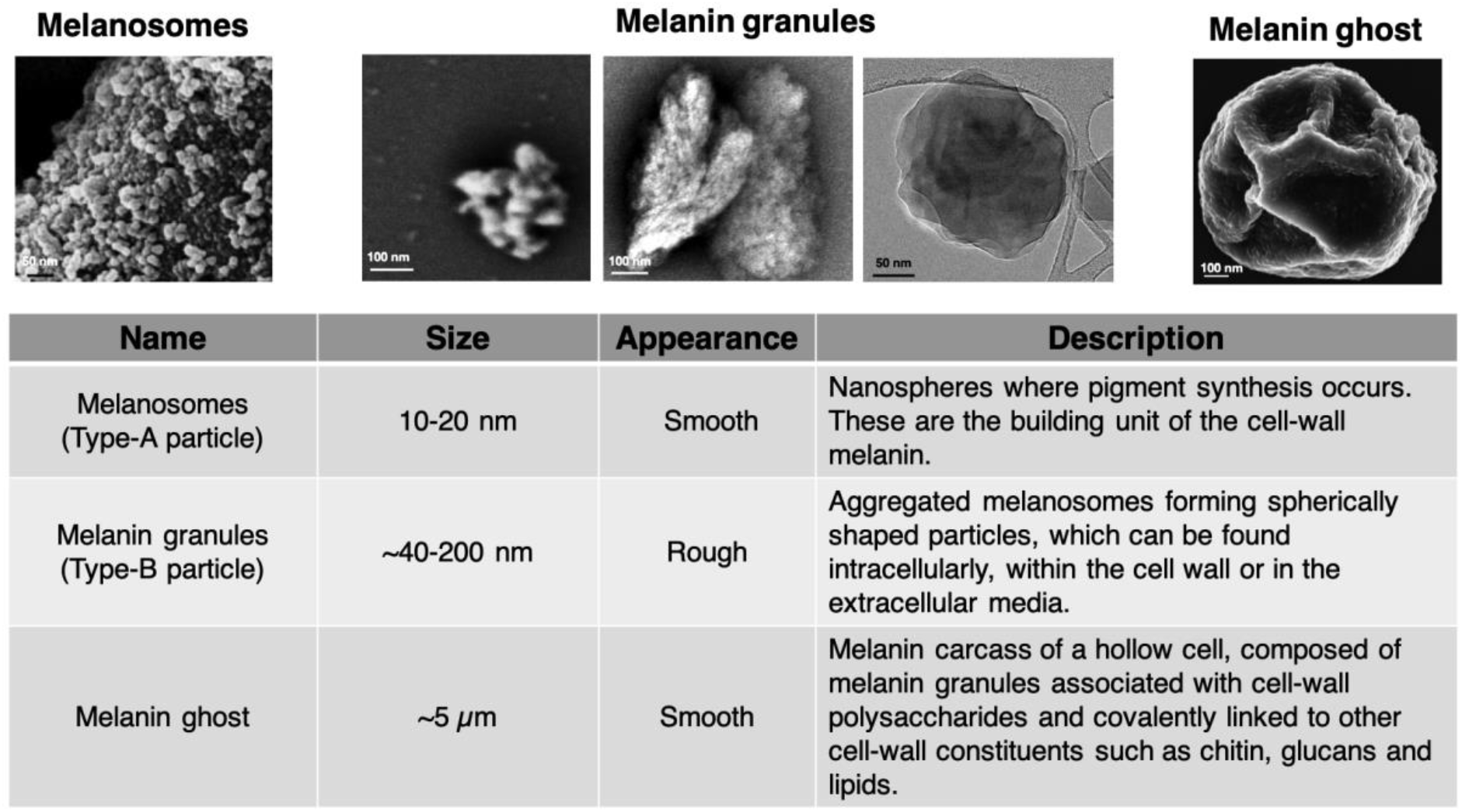
Summary of *C. neoformans* melanin particles visualized with different microscopic approaches.

## Discussion

Elucidation of the smallest fungal cell-wall melanin particle that retains its characteristics from *C. neoformans* is essential to understanding how the pigment is assembled and interacts with other cellular components. Similarly, an appreciation of melanin supramolecular structure (40) is essential to understand how this material confers upon the fungal cell with unique biological properties beyond microbial virulence, such as photoprotection, energy harvesting, heat gain, thermoregulation, and metal scavenging, among others (3).

Previous structural studies on the molecular architecture of fungal eumelanins in the cell wall have focused on the so-called melanin “ghosts”, which are macromolecular structures that retain cellular morphology after hot acid digestion of melanized cells (16,27,49). Although such studies have been extremely informative in revealing chemical aspects of melanin composition, they focus on structures where the pigment remains organized into a cellular carcass of what was once a melanized cell. As such, this melanin polymer includes non-melanin components that serve to hold the pigment together into structures with cell-like dimensions. In this study, we sought to go deeper in searching for the building block unit of the cell-wall melanin through complementary approaches that included prolonged acid digestion of melanin ghosts as well as to investigate the presence of such structures in culture supernatants. Both approaches produced much smaller melanin particles that were comparable in size and form. We then applied multiple techniques to study these particles: SEM, TEM, and cryoEM were used to examine the surface and cross-sectional images of the melanin nanoparticles. DLS was used to analyze the size distribution of melanin particles in suspension and to probe their colloidal properties. Isopycnic centrifugation was used to isolate melanin granules based on their buoyancy. Spectroscopic methodologies (optical absorbance, EPR, and ssNMR) were used to probe the molecular properties of fungal eumelanins and investigate their interaction with cell-wall components. Lastly, proteomic analysis was conducted to identify proteins that maintained an intimate association with melanin granules. The results demonstrate the existence of relatively monodisperse melanin granules ranging from 40 to 200 nm in diameter, depending on the approach, that manifested a monotonic broadband absorption curve and stable EPR signal consistent with melanin. These particles are in turn composed of melanosomes, substructural nanospheres serving as the building unit of the fungal cell wall, functionally comparable to those described in mammalian melanization systems (66).

The small melanin particles derived from “ghosts” or culture supernatants are different from those obtained by auto-polymerization of L-DOPA into melanin, implying differences between the products of the biological catalytic and chemical reactions. Melanin “ghosts” can be broken down to smaller particles by either acid digestion or sonic cavitation generating particles that are comparable in size to those released into the culture supernatant during *C. neoformans* growth under melanizing conditions. The differences between *C. neoformans* derived particles and those from auto-polymerized L-DOPA imply different synthesis, as fungal particles showed a unique electron microscopic appearance and size. We have shown previously that *C. neoformans* and synthetic melanins shared similar chemical groups in the aromatic core (18); however, differences between these materials could stem from the fact that the production of fungal particles is vesicle-associated and catalyzed by a laccase, whereas auto-polymerization is a spontaneous oxidative process that occurs in solution. Moreover, *C. neoformans* melanin granules isolated from the extracellular media are spherical shaped and biologically synthesized particles of up to 200 nm in diameter, similar to those isolated from natural eumelanins such as *Sepia officinales* (43,67), and are found covalently bound to proteins as also reported for mammalian melanin (68).

The fungal melanin granules demonstrated a close association with four proteins relevant to *C. neoformans* virulence: Cig1, Blp1, Qsp1, and CNI3590 macrophage-activating glycoprotein (**Table 1**). The mannoprotein Cig1 (CNAG_01653) plays a role in iron acquisition and it is essential for virulence in a mutant lacking the ferroxidase for high-affinity uptake (69); no defect in melanin production is reported for the *C. neoformans cig1*Δ mutant (61). The hypothetical protein containing a Barwin-like domain (Blp1) (CNAG_06346) is involved in capsule-independent anti-phagocytosis activity and is critical for successful colonization of mammalian lungs by *C. neoformans*. Blp1 is not essential for capsule formation and melanin production (63), but this domain is reported to show binding affinity for chitin oligomers (70), suggesting a possible association of this protein with the cell-wall chitin. The quorum sensing peptide (Qsp1) (CNAG_03012) is a secreted central signaling molecule that acts intracellularly to control virulence in *C. neoformans* and promote cell wall function at high cell densities (64,71). *QSP1* mutant strains display hypomelanization at 37°C but no capsule reduction (64). The melanization process requires the intracellular released of the mature Qsp1 peptide (64), which could explain the finding of this unique sequence in our proteomic analyses of melanin granules. CNI3590 (CNAG_05312) is a GPI-anchored protein and also a secreted factor associated with virulence that contains a carbohydrate-binding domain (62,63), suggesting its interaction with the plasma membrane and cell-wall components such as glucans, chitin and chitosan. Furthermore, Cig1 and CNAG_05312 are positively regulated by the pH-responsive transcription factor Rim101, which in turns is induced by Pka1, a catalytic subunit of the cyclic-AMP/Protein Kinase A (cAMP/PKA) pathway (62,72). Pka1 is known to modulate the secretion of proteins associated with virulence and survival of *C. neoformans* in the host, as well as the expression of cell-wall related genes (62,73). Similarly, Blp1, Qsp1 and CNAG_05312 are direct targets of three transcription factors required for *C. neoformans* virulence: Gat201, Gat204 and Liv3 (63,64,74). It is noteworthy that *pka*Δ mutant strains fail to produce melanin or capsule (75), whereas deletion of *GAT201* reduces the capsule size and severely attenuates virulence in a murine model but shows a hypermelanized colony phenotype (74). In contrast, neither *gat204*Δ (63) nor *liv3*Δ (74) mutant strains exhibit defects in melanin or capsule production. Altogether, these data strongly support the notion that *C. neoformans* synthesis of melanin granules is a highly regulated process that leads to particles with unique morphological and compositional features, whose biogenesis might be subject to highly redundant gene networks. Although we cannot rule out that these proteins become associated with the melanin particles in culture supernatants, covalent association of melanin to proteins has been extensively demonstrated in mammalian melanogenesis (68). The association between *C. neoformans* melanin granules and these proteins may serve complex regulatory roles during the biopolymer synthesis and structural functions within the granular particle or the fungal cell wall milieu.

*C. neoformans* melanin granules visualized by negative staining TEM and cryoEM most likely are formed from the aggregation of melanosomes (∼30 nm in diameter), resembling the supramolecular structure described in *Sepia* melanin (43) and biomimetic eumelanin (46). Analysis of particles resulting from eumelanin synthesis via enzyme immobilization prompted the proposal of a mechanism of hierarchical buildup of melanin by the gradual aggregation of building block particles (type-A particles), which led to the formation of particles 200 nm in diameter (type-B particles) (46). Prior studies from our group noted that melanin ghosts are composed of smaller melanin granules arranged in concentric layers embedded within the fungal cell wall (49). Ghost structures revealed that the smallest melanin nanospheres with smooth appearance are seen on the surface of early buds. We propose that as the fungal cell ages, aggregated melanosomes giving rise to melanin granules (up to 200 nm in diameter) are embedded into the cell wall. These melanin particles display a layered concentric arrangement guided by membranous sheets within the cryptococcal cell wall, which becomes evident by the removal of non-pigment acid-labile cell-wall constituents. This supramolecular structure is also consistent with the findings of previous studies using NMR cryoporometry, which indicated that *C. neoformans* melanin layers have pores that become smaller with increased cell age and progressive melanization (49).

Moreover, our micrographs showed a close association between “finger-like” plasma membrane invaginations and melanin granules, suggesting that these membranes structures could be involved in transport from the cytosol to the fungal cell wall. In *Saccharomyces cerevisiae*, “finger-like” invaginations are sphingolipid-rich lateral domains termed MCP (membrane compartment of Pma1) (76–78). These domains are characterized by cortical patches of fibrous actin involved with vesicular traffic (79), most likely associated with an “inverted macropinocytosis” mechanism required for extracellular vesicles transport across the plasma membrane and into the cell wall (80). Previous vesicles studies in *C. neoformans* have reported invaginations of the plasma membrane (81) and revealed an intimate association with sphingolipids and sterols (26). The exact nature of lateral membrane microdomains involved in the vesicular transport of *C. neoformans* and the role of lipids in this context remains to be elucidated.

Mammalian melanin synthesis is enzymatically driven and occurs in membrane-enclosed compartments called melanosomes, which prevents the exposure of most cellular components to toxic or deleterious reactants produced during the process (82). These specialized intracellular organelles are characterized by different developmental stages (I-IV) (25) that progressively incorporate pigment cell-specific components such as Pmel17 (83), “lipid rafts” critical for sorting of tyrosinase and other melanogenic proteins (84). Structurally, eumelanosomes (most pigment content corresponds to eumelanin) are aggregated structures (type-B particles) of 95-307 nm in diameter, depending on the source, formed due to the previous assembly of 10-60 nm subunits of small nanoparticles (type-A particles) (45). Analogous to this system, fungal melanogenesis is also compartmentalized within a sphingolipid-enriched vesicle, depends on the laccase enzyme, and leads to the supramolecular buildup of cryptococcal eumelanin particles that are retained within the cell wall, where they provide protection against environmental stressors (3,85,86) and interfere with the host immune responses (20,87).

Our proposed model for melanization in *C. neoformans* is in consonance with previous observations by our group and others. We have identified organelles showing that pigment synthesis starts intracellularly within melanosomes, enclosed in double-layered compartments, which are then transferred to the cell wall to consolidate as melanin granules anchored within the cell wall. Several pieces of evidence are presented, which can be organized into a testable model. First, laccase is expressed within MVBs as well as in the cell wall (88,89). *C. neoformans* melanized vesicles measuring between 320 to 380 nm in diameter are shown to incorporate L-DOPA as an electron-dense material (27), suggesting an explanation to the increased number of multivesicular protrusions seen in melanized versus non-melanized cells (60) (**Figure 12A**). Next, aggregated melanosomes ∼60 nm in diameter (**Figure 12B, lower inset**) may rely on MCP-like regions of the plasma membrane that generate finger-like protrusions that mediate melanosomes’ traffic to the cell wall (**Figures 12C,D**). A previous study revealed that the major capsular component (GXM, glucoronoxylomannan) was found in association with endosome-like membranous structures possibly involved in GXM synthesis/transport (90). The plasma membrane is a dynamic organelle compartmentalized into two lateral microdomains with specific functions and composition (91). One of these highly conserved domains in *S. cerevisi*ae is closely related to an unconventional mechanism for vesicle secretion termed inverted macropinocytosis (79,80). Lastly, ongoing aggregation of melanin granules within the cell wall is supported by eumelanin biomimetic studies demonstrating that consolidation of type-B particles is due to high amounts of the laccase enzyme (46), which is in agreement with increased laccase localization at the outer edge of the cell wall versus the inner edge (92). Melanin layered deposition on the cell wall follows the existence of membranous sheets described in early ultrastructural studies of the cryptococcal cell wall (93,94) (**Figure 12F**); however, as we have demonstrated in this study melanin particles are loosely associated with an acid-labile polysaccharide matrix that makes this layout within the melanized cell wall difficult to demonstrate. Nonetheless, the concentric layered arrangement of these granular particles ∼75 nm in diameter was exposed when analyzing melanin ghosts (49,95). The covalent binding of melanin to aliphatic moieties (polysaccharides, chitin/chitosan and lipids) has been shown by ss-NMR analyses (16,19), possibly favored via electrostatic interactions between chitin/chitosan and melanin. Indeed, alterations in the cell-wall chitin/chitosan induced a “leaky melanin” phenotype (31,32,34) or an enhanced deposition of the pigment (14). The recovery of melanin granules in the supernatant of the culture media is explained by normal cell-wall remodeling required for cellular budding (**Figures 5A,B; 9A,B**), which involves local modifications of the melanin scaffold by secreted peptidases (96), chitinases (97), and glucanases (62) that lead to melanin granules detachment from the cell wall. The model does not address the source of strongly associated lipids with melanin granules, but rather provides new pieces of data that should encourage experiments to continue dissecting the phenomenon of fungal cell wall melanization.

In summary, we note remarkable parallels between cryptococcal and mammalian melanization. We have assembled the available facts into a model for the cryptococcal melanogenesis process that can be tested in future studies. The resemblances in these two systems, are perhaps not surprising given that animals and fungi are close relatives in the tree of life and that the chemistry of melanin synthesis is similar. In both systems, the melanization reaction is contained within a vesicular structure, which protects the host cell from the cytotoxicity of reaction intermediates. The isolation of melanin particles from culture supernatants promises to be a new source of melanin material for studies that further unlock the details of pigment formation and for raw materials in industrial applications include radioprotection, light harvesting, and drug delivery, among various uses.

## Experimental procedures

### Strains

In this study, the following strains were used: *C. neoformans* strain H99 (serotype A), *C. neoformans* Δ*LAC1,2* (H99 background), and *C. neoformans* ST211A (H99 background) kindly provided by Alexander Idnurm. Each strain was kept in 20% glycerol stocks at −80°C, and thawed on ice when needed for replication.

### Cell growth and culture conditions

For fungal culture growth, a loop of the glycerol stock was inoculated into Sabouraud broth and incubated at 30 °C for 48 h. Experiments were performed using cultures grown for 10-14 days at 30 °C with continuous shaking at 120 rpm in defined chemical media [minimal media (MM): 15 mM dextrose, 10 mM MgSO_4_, 29.4 mM KH_2_PO_4_, 13 mM glycine, 3 μM thiamine-HCl, pH 5.5] with and without 1 mM L-DOPA.

### Extended acid-hydrolysis of melanin ghosts

Melanin ghosts were isolated following the protocol previously described by Chatterjee *et al* (16). Briefly, cells grown in MM with L-DOPA were collected by centrifugation and washed twice with phosphate buffer (pH 7.4). Melanized cells were then subjected to a multiple-step enzymatic and chemical treatment to remove cell-wall, proteins, and lipid components associated with the melanin polymer. Consequently, the end product is a hollowed particle coated by the acid-resistant pigment called a “melanin ghost” (59). The acid treatment step consists of incubation of the melanized particles in 6 N HCl at 80°C for 1 h. For the extended acid-hydrolysis, melanin ghosts were incubated for up to 4 days including periodic addition of fresh acid solution. Aliquots at 1 h, 24 h and 4 d were analyzed by scanning electron microscopy (SEM). The supernatant from the acid-hydrolysis was also collected, visualized by transmission electron microscopy (TEM) and SEM. Its particle population size was estimated by measuring the diameter using SEM micrographs.

### Transmission Electron Microscopy (TEM)

*C. neoformans* yeast cells or melanin particles were fixed in 2.5% (v/v) glutaraldehyde in 0.1 M sodium phosphate buffer (pH 7.3) for 24 h at 4°C. Samples were encapsulated in 3% (w/v) low-melting-point agarose prior to being processed in Spurr resin following a 24-h schedule on a Lynx tissue processor (secondary 1% osmium tetroxide fixation; 1% uranyl acetate contrasting; ethanol dehydration and infiltration with acetone/Spurr resin). Additional infiltration was conducted under vacuum at 60°C before the samples were embedded in TAAB capsules (TAAB Laboratories, Amersham, United Kingdom) and polymerized at 60°C for 48 h. Semithin survey sections 0.5 μM thick were stained with 1% toluidine blue to identify the areas with the best cell density. Ultrathin sections (60 to 90 nm) were prepared with a Diatome diamond knife on a Leica UC6 ultramicrotome and stained with 2% uranyl acetate in 50% methanol, followed by lead citrate for examination with a JEOL 1200EX (Analytical Imaging Facility at Albert Einstein College of Medicine) or a Philips CM120 at 80 kV (Microscopy Facility at the Johns Hopkins School of Medicine). Negative staining of the fractions from OptiPrep density gradients was performed by adsorbing 10 μL of each fraction to glow-discharged 400 mesh ultra-thin carbon coated grids (EMS CF400-CU-UL) for two minutes, followed by 3 quick rinses of TBS and staining with 1% uranyl acetate with 0.05% Tylose. Images were captured with an Gatan Orlus Digital 2K x 2K CCD camera or an AMT XR80 high-resolution (16-bit) 8 Megapixel camera.

### Scanning electron microscopy (SEM)

Melanin particles were suspended in PBS and incubated on coverslips coated with 0.01% polylysine and fixed with 2.5% glutaraldehyde in 0.1 M sodium phosphate buffer (pH 7.3) for 24 h at 4°C. Cells and melanin particles were postfixed with 1% OsO_4_; dehydrated in 70%, 90%, 95%, and 100% ethanol; and critical-point dried in CO_2_. The cells were sputter coated with gold using an EMitech K550 Sputter Coater and viewed in a LEO/Zeiss Field-emission scanning electron microscope at a voltage of 10 kV.

### Dynamic Light Scattering (DLS)

Dynamic light scattering (DLS) techniques gives an estimate of the size and heterogeneity of a sample by measuring the intensity fluctuations of scattered light by particles in solution. Measurement of melanin particles by DLS was performed with a 90Plus instrument (Brookhaven Instruments). To assess the size of melanin ghosts after different sonication times, five hundred microliters of a melanin ghost suspension in distilled water (1 mg/ml) was sonicated with a horned sonicator at amplitude 5 for up to 12 minutes. Particle size of the suspension was measured at different time points. Data are expressed as the average of 10 runs of 1 min data collection each. To examine the colloidal properties as a function of salt concentration, 1M, 0.1 M, 0.01 M, and 0.001 M solutions of Sodium Chloride (NaCl) and Calcium Chloride (CaCl2) and 10X, 1X, 0.1X and 0.01X Phosphate Buffered Saline solutions were used to suspend 1 μl of vesicles/granules in 100 μl of the respective salt solution. Average hydrodynamic diameter was obtained as above mentioned.

### Isolation of melanin granules

The isolation of melanin granules from *C. neoformans* was performed in two phases: 1) Isolation a mixture of extracellular vesicles and melanin granules from the culture supernatant herein called crude melanin granules, and 2) Isolation of melanin granules by density gradient. For the first phase, yeast cells were pelleted at 5,000 rpm (SLA rotor, Sorvall) for 30 min at 4°C. The supernatant was filtered through a 0.22 μm pore filter (Millipore Sigma, USA) and pelleted at 100,000 x *g* for 1 hour at 4 °C, with a slow break (SW28 rotor, Beckman Coulter). The pellet was suspended in 1X Dulbecco’s Phosphate-Buffer Saline (DPBS) without calcium and magnesium (Corning, USA) and washed twice with the same buffer. The pellet (crude melanin granules) was suspended in approximately 500 μl of 0.1X DPBS and stored at −80°C until further analysis. For the second phase, OptiPrep™ solution (60%) dilutions were made in 10 mM HEPES, 0.85% NaCl at pH 7.4. The crude melanin granules were mixed with undiluted OptiPrep™ to make a 45% solution that was layered at the bottom of the gradient. The gradient was comprised of 45%, 35%, 30%, 25%, 20% and 15% layers in the ratio 0.4:3:3:2:2:1 for a total volume of 3 ml. The gradient was spun at 50,000 rpm for 10 hours (SW Ti55 rotor, Beckman Coulter) to allow the vesicles and granules to equilibrate at their respective densities. Five equal volume fractions were collected from each tube.

### Spectroscopic studies

#### UV-Visible analysis

The absorption properties of crude vesicle samples were analyzed with UV-Vis spectroscopy using a SpectraMax M2 microplate reader (Molecular Devices, USA). The spectra were recorded in the wavelength range of 300 to 800 nm.

#### Electron Paramagnetic Resonance (EPR)

Suspensions of melanin samples (∼2 mg) in 150 μl distilled water were sonicated for 1 min using a horned sonicator. Equal volumes of appropriated buffer (distilled water at pH 2.0, 7.0, 12.0 or 0.1 M ZnCl_2_) was added. Using a glass long-tip transfer pipette, each sample was placed in a 4-mm precision-bore quartz EPR tubes and frozen by immersion in liquid nitrogen. The EPR spectra were recorded at 77 K using a Bruker EMX EPR spectrometer operating in X-band mode. Experimental parameters were as follows: modulation amplitude, 10 G; microwave power, 6.362 mW; modulation frequency, 100 kHz; microwave frequency, 9.43 GHz; attenuation, 15 dB; number of scans averaged, 20. Data acquisition was performed using WinEPR software (Bruker). All spectra were obtained under identical instrumental conditions. Light irradiation of samples was carried out at room temperature for 20 min using a 250 W cool white light operating at 4000K.

#### Solid-State Nuclear Magnetic Resonance (ss-NMR)

Solid-state NMR experiments were carried out on a 600 MHz Varian (Agilent) DirectDrive2 (DD2) spectrometer using a 1.6 mm FastMAS probe (Agilent Technologies, Santa Clara, CA, USA) with a magic-angle spinning (MAS) rate of 15.00±0.02 kHz and a nominal spectrometer-set temperature of 25 °C. All spectra were acquired on ^13^C-enriched samples obtained from a *C. neoformans* ST211A cell culture containing [U-^13^C_6_-glucose] and natural abundance L-DOPA as the sole carbon source and obligatory melanization precursor, respectively. Average post-lyophilization sample masses post lyophilization were ∼1-3 mg for the pelleted cellular material containing melanin granules and ∼5-7 mg for the melanin ghosts. The typical 90 degree pulse_lengths for the 13C cross-polarization (CPMAS) experiments were 1.65 μs for ^1^H and 2.25 μs for ^13^C, respectively. As described previously (18), ^1^H decoupling using the small phase incremental alternation pulse sequence (SPINAL) with a field strength of ∼180 kHz was applied during signal acquisition. The 2D ^13^C-^13^C through-space dipolar assisted rotational resonance (DARR) correlation experiments were collected using a 50 ms mixing time with similar ^1^H and ^13^C 90 degree pulse-lengths and decoupling field strength as described for the ^13^C CPMAS experiments.

### CryoEM

Five microliters of purified *C. neoformans* melanin granules (∼3 mg) was applied to a 300 mesh copper grid coated with a holey carbon film; and vitreously frozen in liquid nitrogen-cooled ethane using an FEI Vitrobot Mark IV device. The sample was transferred to a Gatan 626/70° cryo-transfer holder; and observed in an FEI Talos S200C transmission electron microscope operating at 200 kV. Images were collected with a Falcon 2 direct-electron detector camera; and processed in Adobe Photoshop CS6 with only linear adjustments in brightness and contrast.

### Proteomic analysis using mass spectrometry

*C. neoformans* isolated melanin granules and the equivalent fraction from non-melanized material (∼3 mg of each sample) were treated following a protocol described previously (98). Briefly, samples were subjected to in-solution digestion by resuspending in a solubilization buffer [8 M urea, 10% acetonitrile in 100 mM ammonium bicarbonate (pH 8.5)], flash frozen and thawed in a water bath at 30 °C for three rounds. Each sample was sonicated briefly on ice (10 s, horned sonicator at maximum amplitude) and mixed using a vortexer. Next, samples were reduced with 10 mM dithiothreitol (DTT) (Sigma) at 51°C for 1 h and carboxyamido-methylated with 20 mM iodoacetamide (Sigma) for 45 min in the dark at room temperature. Subsequent digestions with Endo-Protease Lys-C (Roche Diagnostics, USA) and trypsin (1:20 enzyme/protein ratio) for 6 h and 10 h, respectively, were performed at room temperature. The reaction was stopped via addition of 10% trifluoroacetic acid (TFA) and incubation for 30 min at 37 °C. Lastly, samples were centrifuged for 15 min at 20,800 g, supernatants were individually transferred to low-protein binding 1.5 ml tubes (Eppendorf) and completely dried in acid-resistant CentriVap vacuum concentrator (Labconco). Peptides were resuspended in 40 μl of 0.1% TFA and stored at −80°C until the mass spectrometric analysis could be performed. Digested samples were cleaned up on an Oasis HLB micro-elution plate and evaporated to dryness. The samples were taken up in 2% acetonitrile, 0.1% formic acid then vortexed and centrifuged before placing into a 96 well plate. The samples were injected using an EasyLC into a QE-Plus (Thermo Scientific) mass spectrometer and eluted over a 90 minute gradient from 100% buffer A (2% acetonitrile, 0.1% formic acid) to 100% buffer B (90% acetonitrile, 0.1% formic acid) with an MS resolution of 70,000 and an MS2 resolution of 35,000 running a top 15 DDA method. Target values for MS and MS2 were set at 3e6 and 1e5 with 100 and 150 milliseconds, respectively and a normalized collision energy of 28. Data were searched against the *Cryptococcus neoformans* database (Broad Institute, November 2016) using PEAKS version 7 (Bioinformatics Solutions Inc., Waterloo, ON, Canada) using a peptide tolerance of 5 ppm and a fragment ion tolerance of 0.02 daltons. Enzyme specificity was set to semi specific trypsin with 2 missed cleavages allowed and variable carbamidomethyl C, oxidation M, and deamidation N,Q. Search results were reported at a 1% FDR for all peptides.

### *In silico* predictions

Bioinformatic analyses to identify the presence of signal peptide and glycophosphatidylinositol (GPI) anchor were performed using SignalP v. 5.0 (99) and GPI-SOM (100), respectively.

## Acknowledgements

*We are thankful to Alexander Idnurm for kindly providing us with C. neoformans strain ST211A, Carolina Coelho for critical reading of the manuscript, Andre Nicola and Alexander Alanio for valuable discussion of experimental designs. We also gratefully acknowledge expertise and technical support from Barbara Smith (Microscopy Facility, School of Medicine, Johns Hopkins University) with transmission EM, SEM and negative staining. We thank Joel Tang for technical support for data acquisition in connection with EPR analyses. This research was supported by a grant from the U.S. National Institutes of Health (NIH R01-AI052733). The NMR facilities used in this work are operated by The City College and the CUNY Institute for Macromolecular Assemblies, with additional infrastructural support provided by NIH 8G12 MD007603 from the National Institute on Minority Health and Health Disparities of the National Institutes of Health. E.C. was the recipient of a Johns Hopkins Malaria Research Institute fellowship. C.C. was the recipient of a fellowship award from the U.S. Department of Education Graduate Assistance in Areas of National Need (GAANN) Program in Biochemistry, Biophysics, and Biodesign at The City College of New York (PA200A120211 and PA200A150068). RP-R was supported in part by NIH/NIAID grant AI115091. RP-R was further a ‘Ramon y Cajal’ fellow from the Spanish Ministry of Economy and Competitiveness. RP-R was also supported by the Spanish Ministry of Economy and Competitiveness [grant SAF2016-77433-R]. RP-R acknowledges support from CICbioGUNE through the Severo Ochoa Excellence Accreditation (SEV-2016-0644). JMMc wishes to acknowledge the support of the Hopkins Integrated Imaging Center; and NIH grant* NIH-NCRR 1S10OD012342 for the acquisition of the Talos TEM.

## References

1. Shosuke, I., Kazumasa, W., Marco, d. I., Alessandra, N., and Alessandro, P. (2011) Structure of Melanins. in Melanins and Melanosomes: Biosynthesis, Biogenesis, Physiological, and Pathological Functions. (Borovansky, J., and Patrick A., R. eds.), First Ed., Wiley-VCH verlag & Co. KGaA, Germany. pp 167–185

2. Hill, H. Z. (1992) The function of melanin or six blind people examine an elephant. Bioessays 14, 49–56

3. Cordero, R. J. B., and Casadevall, A. (2017) Functions of fungal melanin beyond virulence. Fungal Biology Reviews 31, 99–112

4. Christensen, B. M., Li, J., Chen, C. C., and Nappi, A. J. (2005) Melanization immune responses in mosquito vectors. Trends Parasitol 21, 192–199

5. White, L. P. (1958) Melanin: a naturally occurring cation exchange material. Nature 182, 1427–1428

6. Gomez, B. L., and Nosanchuk, J. D. (2003) Melanin and fungi. Curr Opin Infect Dis 16, 91–96

7. Huffnagle, G. B., Chen, G. H., Curtis, J. L., McDonald, R. A., Strieter, R. M., and Toews, G. B. (1995) Down-regulation of the afferent phase of T cell-mediated pulmonary inflammation and immunity by a high melanin-producing strain of Cryptococcus neoformans. J Immunol 155, 3507–3516

8. Wang, Y., Aisen, P., and Casadevall, A. (1995) Cryptococcus neoformans melanin and virulence: mechanism of action. Infect Immun 63, 3131–3136

9. Wang, Y., and Casadevall, A. (1994) Growth of Cryptococcus neoformans in presence of L-dopa decreases its susceptibility to amphotericin B. Antimicrob Agents Chemother 38, 2648–2650

10. Tajima, K., Yamanaka, D., Ishibashi, K.-i., Adachi, Y., and Ohno, N. (2019) Solubilized melanin suppresses macrophage function. FEBS Open Bio, 1–10

11. Chaskes, S., and Tyndall, R. L. (1975) Pigment production by Cryptococcus neoformans from para- and ortho-Diphenols: effect of the nitrogen source. J Clin Microbiol 1, 509–514

12. Chaskes, S., and Tyndall, R. L. (1978) Pigment production by Cryptococcus neoformans and other Cryptococcus species from aminophenols and diaminobenzenes. J Clin Microbiol 7, 146–152

13. Kwon-Chung, K. J., Tom, W. K., and Costa, J. L. (1983) Utilization of indole compounds by Cryptococcus neoformans to produce a melanin-like pigment. J Clin Microbiol 18, 1419–1421

14. Camacho, E., Chrissian, C., Cordero, R. J. B., Liporagi-Lopes, L., Stark, R. E., and Casadevall, A. (2017) N-acetylglucosamine affects Cryptococcus neoformans cell-wall composition and melanin architecture. Microbiology 163, 1540–1556

15. Chatterjee, S., Prados-Rosales, R., Frases, S., Itin, B., Casadevall, A., and Stark, R. E. (2012) Using solid-state NMR to monitor the molecular consequences of Cryptococcus neoformans melanization with different catecholamine precursors. Biochemistry 51, 6080–6088

16. Chatterjee, S., Prados-Rosales, R., Itin, B., Casadevall, A., and Stark, R. E. (2015) Solid-state NMR Reveals the Carbon-based Molecular Architecture of Cryptococcus neoformans Fungal Eumelanins in the Cell Wall. J Biol Chem 290, 13779–13790

17. Chatterjee, S., Prados-Rosales, R., Tan, S., Itin, B., Casadevall, A., and Stark, R. E. (2014) Demonstration of a common indole-based aromatic core in natural and synthetic eumelanins by solid-state NMR. Org Biomol Chem 12, 6730–6736

18. Chatterjee, S., Prados-Rosales, R., Tan, S., Phan, V. C., Chrissian, C., Itin, B., Wang, H., Khajo, A., Magliozzo, R. S., Casadevall, A., and Stark, R. E. (2018) The melanization road more traveled by: precursor substrate effects on melanin synthesis in cell-free and fungal cell systems. J Biol Chem

19. Zhong, J., Frases, S., Wang, H., Casadevall, A., and Stark, R. E. (2008) Following fungal melanin biosynthesis with solid-state NMR: biopolymer molecular structures and possible connections to cell-wall polysaccharides. Biochemistry 47, 4701–4710

20. Nosanchuk, J. D., and Casadevall, A. (2003) The contribution of melanin to microbial pathogenesis. Cell Microbiol 5, 203–223

21. Eisenman, H. C., and Casadevall, A. (2012) Synthesis and assembly of fungal melanin. Appl Microbiol Biotechnol 93, 931–940

22. Hegnauer, H., Nyhlen, L., and Rast, D. (1985) Ultrastructure of native and synthetic Agaricus bisporus melanins. Implications as to the compartmentation of melanogenesis in fungi. Exp Mycol 9, 221–229

23. Alviano, C. S., Farbiarz, S. R., De Souza, W., Angluster, J., and Travassos, L. R. (1991) Characterization of Fonsecaea pedrosoi melanin. J Gen Microbiol 137, 837–844

24. San-Blas, G., Guanipa, O., Moreno, B., Pekerar, S., and San-Blas, F. (1996) Cladosporium carrionii and Hormoconis resinae (C. resinae): cell wall and melanin studies. Curr Microbiol 32, 11–16

25. Seiji, M., Fitzpatrick, T. B., Simpson, R. T., and Birbeck, M. S. (1963) Chemical composition and terminology of specialized organelles (melanosomes and melanin granules) in mammalian melanocytes. Nature 197, 1082–1084

26. Rodrigues, M. L., Nimrichter, L., Oliveira, D. L., Frases, S., Miranda, K., Zaragoza, O., Alvarez, M., Nakouzi, A., Feldmesser, M., and Casadevall, A. (2007) Vesicular polysaccharide export in Cryptococcus neoformans is a eukaryotic solution to the problem of fungal trans-cell wall transport. Eukaryot Cell 6, 48–59

27. Eisenman, H. C., Frases, S., Nicola, A. M., Rodrigues, M. L., and Casadevall, A. (2009) Vesicle-associated melanization in Cryptococcus neoformans. Microbiology 155, 3860–3867

28. Franzen, A. J., Cunha, M. M., Miranda, K., Hentschel, J., Plattner, H., da Silva, M. B., Salgado, C. G., de Souza, W., and Rozental, S. (2008) Ultrastructural characterization of melanosomes of the human pathogenic fungus Fonsecaea pedrosoi. J Struct Biol 162, 75–84

29. Walker, C. A., Gomez, B. L., Mora-Montes, H. M., Mackenzie, K. S., Munro, C. A., Brown, A. J., Gow, N. A., Kibbler, C. C., and Odds, F. C. (2010) Melanin externalization in Candida albicans depends on cell wall chitin structures. Eukaryot Cell 9, 1329–1342

30. Upadhyay, S., Xu, X., Lowry, D., Jackson, J. C., Roberson, R. W., and Lin, X. (2016) Subcellular Compartmentalization and Trafficking of the Biosynthetic Machinery for Fungal Melanin. Cell Rep 14, 2511–2518

31. Baker, L. G., Specht, C. A., Donlin, M. J., and Lodge, J. K. (2007) Chitosan, the deacetylated form of chitin, is necessary for cell wall integrity in Cryptococcus neoformans. Eukaryot Cell 6, 855–867

32. Banks, I. R., Specht, C. A., Donlin, M. J., Gerik, K. J., Levitz, S. M., and Lodge, J. K. (2005) A chitin synthase and its regulator protein are critical for chitosan production and growth of the fungal pathogen Cryptococcus neoformans. Eukaryot Cell 4, 1902–1912

33. Walton, F. J., Idnurm, A., and Heitman, J. (2005) Novel gene functions required for melanization of the human pathogen Cryptococcus neoformans. Mol Microbiol 57, 1381–1396

34. Tsirilakis, K., Kim, C., Vicencio, A. G., Andrade, C., Casadevall, A., and Goldman, D. L. (2012) Methylxanthine inhibit fungal chitinases and exhibit antifungal activity. Mycopathologia 173, 83–91

35. Bull, A. T. (1970) Chemical composition of wild-type and mutant Aspergillus nidulans cell walls. The nature of polysaccharide and melanin constituents. J Gen Microbiol 63, 75–94

36. Wang, Z., Zheng, L., Hauser, M., Becker, J. M., and Szaniszlo, P. J. (1999) WdChs4p, a homolog of chitin synthase 3 in Saccharomyces cerevisiae, alone cannot support growth of Wangiella (Exophiala) dermatitidis at the temperature of infection. Infect Immun 67, 6619–6630

37. Cédric, D., Francesca, G., Michael S, M., and Graça, R. (2011) Biogenesis of melansomes. in Melanis and Melanosomes: Biosynthesis, Biogenesis, Physiological, and Pathological Functions (Jan, B., and Patrick A., R. eds.), First Ed., Wiley-VCH Verlag GmbH & Co. KGaA, Germany. pp 247–294

38. d’Ischia, M., Napolitano, A., Pezzella, A., Meredith, P., and Sarna, T. (2009) Chemical and structural diversity in eumelanins: unexplored bio-optoelectronic materials. Angew Chem Int Ed Engl 48, 3914–3921

39. Panzella, L., Ebato, A., Napolitano, A., and Koike, K. (2018) The Late Stages of Melanogenesis: Exploring the Chemical Facets and the Application Opportunities. Int J Mol Sci 19

40. Meredith, P., and Sarna, T. (2006) The physical and chemical properties of eumelanin. Pigment Cell Res 19, 572–594

41. Prota, G. (1988) Progress in the chemistry of melanins and related metabolites. Med Res Rev 8, 525–556

42. Zajac, G. W., Gallas, J. M., Cheng, J., Eisner, M., Moss, S. C., and Alvarado-Swaisgood, A. E. (1994) The fundamental unit of synthetic melanin: a verification by tunneling microscopy of X-ray scattering results. Biochim Biophys Acta 1199, 271–278

43. Clancy, C. M., and Simon, J. D. (2001) Ultrastructural organization of eumelanin from Sepia officinalis measured by atomic force microscopy. Biochemistry 40, 13353–13360

44. Casadevall, A., Nakouzi, A., Crippa, P. R., and Eisner, M. (2012) Fungal melanins differ in planar stacking distances. PLoS One 7, e30299

45. Xiao, M., Chen, W., Li, W., Zhao, J., Hong, Y. L., Nishiyama, Y., Miyoshi, T., Shawkey, M. D., and Dhinojwala, A. (2018) Elucidation of the hierarchical structure of natural eumelanins. J R Soc Interface 15

46. Bungeler, A., Hamisch, B., Huber, K., Bremser, W., and Strube, O. I. (2017) Insight into the Final Step of the Supramolecular Buildup of Eumelanin. Langmuir 33, 6895–6901

47. Ito, S., and Ifpcs. (2003) The IFPCS presidential lecture: a chemist’s view of melanogenesis. Pigment Cell Res 16, 230–236

48. Hong, S., Wang, Y., Park, S. Y., and Lee, H. (2018) Progressive fuzzy cation-pi assembly of biological catecholamines. Sci Adv 4, eaat7457

49. Eisenman, H. C., Nosanchuk, J. D., Webber, J. B., Emerson, R. J., Camesano, T. A., and Casadevall, A. (2005) Microstructure of cell wall-associated melanin in the human pathogenic fungus Cryptococcus neoformans. Biochemistry 44, 3683–3693

50. Nosanchuk, J. D., and Casadevall, A. (2003) Budding of melanized Cryptococcus neoformans in the presence or absence of L-dopa. Microbiology 149, 1945–1951

51. Franzen, A. J., de Souza, W., Farina, M., Alviano, C. S., and Rozental, S. (1999) Morphometric and densitometric study of the biogenesis of electron-dense granules in Fonsecaea pedrosoi. FEMS Microbiol Lett 173, 395–402

52. Walker, L., Sood, P., Lenardon, M. D., Milne, G., Olson, J., Jensen, G., Wolf, J., Casadevall, A., Adler-Moore, J., and Gow, N. A. R. (2018) The Viscoelastic Properties of the Fungal Cell Wall Allow Traffic of AmBisome as Intact Liposome Vesicles. MBio 9

53. Riesz, J., Gilmore, J., and Meredith, P. (2006) Quantitative scattering of melanin solutions. Biophys J 90, 4137–4144

54. Cordero, R. J. B., Robert, V., Cardinali, G., Arinze, E. S., Thon, S. M., and Casadevall, A. (2018) Impact of Yeast Pigmentation on Heat Capture and Latitudinal Distribution. Curr Biol 28, 2657–2664 e2653

55. Nosanchuk, J. D., and Casadevall, A. (1997) Cellular charge of Cryptococcus neoformans: contributions from the capsular polysaccharide, melanin, and monoclonal antibody binding. Infect Immun 65, 1836–1841

56. Momen-Heravi, F., Balaj, L., Alian, S., Mantel, P. Y., Halleck, A. E., Trachtenberg, A. J., Soria, C. E., Oquin, S., Bonebreak, C. M., Saracoglu, E., Skog, J., and Kuo, W. P. (2013) Current methods for the isolation of extracellular vesicles. Biol Chem 394, 1253–1262

57. Sealy, R. C., Hyde, J. S., Felix, C. C., Menon, I. A., and Prota, G. (1982) Eumelanins and pheomelanins: characterization by electron spin resonance spectroscopy. Science 217, 545–547

58. Sarna, T., and Swartz, H. M. (1978) Identification and characterization of melanin in tissues and body fluids. Folia Histochem Cytochem (Krakow) 16, 275–286

59. Wang, Y., Aisen, P., and Casadevall, A. (1996) Melanin, melanin “ghosts,” and melanin composition in Cryptococcus neoformans. Infect Immun 64, 2420–2424

60. Wolf, J. M., Espadas-Moreno, J., Luque-Garcia, J. L., and Casadevall, A. (2014) Interaction of Cryptococcus neoformans extracellular vesicles with the cell wall. Eukaryot Cell 13, 1484–1493

61. Cadieux, B., Lian, T., Hu, G., Wang, J., Biondo, C., Teti, G., Liu, V., Murphy, M. E., Creagh, A. L., and Kronstad, J. W. (2013) The Mannoprotein Cig1 supports iron acquisition from heme and virulence in the pathogenic fungus Cryptococcus neoformans. J Infect Dis 207, 1339–1347

62. Geddes, J. M., Croll, D., Caza, M., Stoynov, N., Foster, L. J., and Kronstad, J. W. (2015) Secretome profiling of Cryptococcus neoformans reveals regulation of a subset of virulence-associated proteins and potential biomarkers by protein kinase A. BMC Microbiol 15, 206

63. Chun, C. D., Brown, J. C. S., and Madhani, H. D. (2011) A major role for capsule-independent phagocytosis-inhibitory mechanisms in mammalian infection by Cryptococcus neoformans. Cell Host Microbe 9, 243–251

64. Homer, C. M., Summers, D. K., Goranov, A. I., Clarke, S. C., Wiesner, D. L., Diedrich, J. K., Moresco, J. J., Toffaletti, D., Upadhya, R., Caradonna, I., Petnic, S., Pessino, V., Cuomo, C. A., Lodge, J. K., Perfect, J., Yates, J. R., 3rd, Nielsen, K., Craik, C. S., and Madhani, H. D. (2016) Intracellular Action of a Secreted Peptide Required for Fungal Virulence. Cell Host Microbe 19, 849–864

65. Hommel, B., Mukaremera, L., Cordero, R. J. B., Coelho, C., Desjardins, C. A., Sturny-Leclere, A., Janbon, G., Perfect, J. R., Fraser, J. A., Casadevall, A., Cuomo, C. A., Dromer, F., Nielsen, K., and Alanio, A. (2018) Titan cells formation in Cryptococcus neoformans is finely tuned by environmental conditions and modulated by positive and negative genetic regulators. PLoS Pathog 14, e1006982

66. Bungeler, A., Hamisch, B., and Strube, O. I. (2017) The Supramolecular Buildup of Eumelanin: Structures, Mechanisms, Controllability. Int J Mol Sci 18

67. Liu, Y., and Simon, J. D. (2003) Isolation and biophysical studies of natural eumelanins: applications of imaging technologies and ultrafast spectroscopy. Pigment Cell Res 16, 606–618

68. d’Ischia, M., Wakamatsu, K., Napolitano, A., Briganti, S., Garcia-Borron, J. C., Kovacs, D., Meredith, P., Pezzella, A., Picardo, M., Sarna, T., Simon, J. D., and Ito, S. (2013) Melanins and melanogenesis: methods, standards, protocols. Pigment Cell Melanoma Res 26, 616–633

69. Jung, W. H., Hu, G., Kuo, W., and Kronstad, J. W. (2009) Role of ferroxidases in iron uptake and virulence of Cryptococcus neoformans. Eukaryot Cell 8, 1511–1520

70. Ludvigsen, S., and Poulsen, F. M. (1992) Three-dimensional structure in solution of barwin, a protein from barley seed. Biochemistry 31, 8783–8789

71. Albuquerque, P., Nicola, A. M., Nieves, E., Paes, H. C., Williamson, P. R., Silva-Pereira, I., and Casadevall, A. (2013) Quorum sensing-mediated, cell density-dependent regulation of growth and virulence in Cryptococcus neoformans. MBio 5, e00986–00913

72. O’Meara, T. R., Norton, D., Price, M. S., Hay, C., Clements, M. F., Nichols, C. B., and Alspaugh, J. A. (2010) Interaction of Cryptococcus neoformans Rim101 and protein kinase A regulates capsule. PLoS Pathog 6, e1000776

73. Hu, G., Steen, B. R., Lian, T., Sham, A. P., Tam, N., Tangen, K. L., and Kronstad, J. W. (2007) Transcriptional regulation by protein kinase A in Cryptococcus neoformans. PLoS Pathog 3, e42

74. Liu, O. W., Chun, C. D., Chow, E. D., Chen, C., Madhani, H. D., and Noble, S. M. (2008) Systematic genetic analysis of virulence in the human fungal pathogen Cryptococcus neoformans. Cell 135, 174–188

75. D’Souza, C. A., Alspaugh, J. A., Yue, C., Harashima, T., Cox, G. M., Perfect, J. R., and Heitman, J. (2001) Cyclic AMP-dependent protein kinase controls virulence of the fungal pathogen Cryptococcus neoformans. Mol Cell Biol 21, 3179–3191

76. Grossmann, G., Opekarova, M., Malinsky, J., Weig-Meckl, I., and Tanner, W. (2007) Membrane potential governs lateral segregation of plasma membrane proteins and lipids in yeast. EMBO J 26, 1–8

77. Mulholland, J., Preuss, D., Moon, A., Wong, A., Drubin, D., and Botstein, D. (1994) Ultrastructure of the yeast actin cytoskeleton and its association with the plasma membrane. J Cell Biol 125, 381–391

78. Gaigg, B., Timischl, B., Corbino, L., and Schneiter, R. (2005) Synthesis of sphingolipids with very long chain fatty acids but not ergosterol is required for routing of newly synthesized plasma membrane ATPase to the cell surface of yeast. J Biol Chem 280, 22515–22522

79. Brach, T., Specht, T., and Kaksonen, M. (2011) Reassessment of the role of plasma membrane domains in the regulation of vesicular traffic in yeast. J Cell Sci 124, 328–337

80. Rodrigues, M. L., Franzen, A. J., Nimrichter, L., and Miranda, K. (2013) Vesicular mechanisms of traffic of fungal molecules to the extracellular space. Curr Opin Microbiol 16, 414–420

81. Rodrigues, M. L., Nakayasu, E. S., Oliveira, D. L., Nimrichter, L., Nosanchuk, J. D., Almeida, I. C., and Casadevall, A. (2008) Extracellular vesicles produced by Cryptococcus neoformans contain protein components associated with virulence. Eukaryot Cell 7, 58–67

82. Delevoye, C., Giordano, F., Marks, M. S., and Raposo, G. (2011) Biogenesis of melanosomes. in Melanins and Melanosomes: Biosynthesis, Biogenesis, Physiological, and Pathological Functions (Borovansky, J., and Riley, P. A. eds.), First Edition Ed., Wiley-VCH Verlag GmbH & Co. KGaA, Weinheim, Germany. pp 247–294

83. Bissig, C., Rochin, L., and van Niel, G. (2016) PMEL Amyloid Fibril Formation: The Bright Steps of Pigmentation. Int J Mol Sci 17

84. Groux-Degroote, S., van Dijk, S. M., Wolthoorn, J., Neumann, S., Theos, A. C., De Maziere, A. M., Klumperman, J., van Meer, G., and Sprong, H. (2008) Glycolipid-dependent sorting of melanosomal from lysosomal membrane proteins by lumenal determinants. Traffic 9, 951–963

85. Martinez, L. R., and Casadevall, A. (2006) Susceptibility of Cryptococcus neoformans biofilms to antifungal agents in vitro. Antimicrob Agents Chemother 50, 1021–1033

86. Steenbergen, J. N., Shuman, H. A., and Casadevall, A. (2001) Cryptococcus neoformans interactions with amoebae suggest an explanation for its virulence and intracellular pathogenic strategy in macrophages. Proc Natl Acad Sci U S A 98, 15245–15250

87. Nosanchuk, J. D., and Casadevall, A. (2006) Impact of melanin on microbial virulence and clinical resistance to antimicrobial compounds. Antimicrob Agents Chemother 50, 3519–3528

88. Zhu, X., Gibbons, J., Garcia-Rivera, J., Casadevall, A., and Williamson, P. R. (2001) Laccase of Cryptococcus neoformans is a cell wall-associated virulence factor. Infect Immun 69, 5589–5596

89. Panepinto, J. C., and Williams, P. R. (2007) The cell biology of virulence - lessons from the pathogenic fungus Cryptococcus neoformans. in Communicating Current Research and Educational Topics and Trends in Applied Microbiology (Mendez-Villas, A. ed.), FORMATEX. pp

90. Oliveira, D. L., Nimrichter, L., Miranda, K., Frases, S., Faull, K. F., Casadevall, A., and Rodrigues, M. L. (2009) Cryptococcus neoformans cryoultramicrotomy and vesicle fractionation reveals an intimate association between membrane lipids and glucuronoxylomannan. Fungal Genet Biol 46, 956–963

91. Foderaro, J. E., Douglas, L. M., and Konopka, J. B. (2017) MCC/Eisosomes Regulate Cell Wall Synthesis and Stress Responses in Fungi. J Fungi (Basel) 3

92. Zhu, X., Gibbons, J., Garcia-Rivera, J., Casadevall, A., and Williamson, P. R. (2001) Laccase of Cryptococcus neoformans Is a Cell Wall-Associated Virulence Factor. Infection and Immunity 69, 5589–5596

93. al-Doory, Y. (1971) The ultrastructure of Cryptococcus neoformans. Sabouraudia 9, 115–118

94. Stoetzner, H., and Kemmer, C. (1971) The morphology of Cryptococcus neoformans in human cryptococcosis. A light-,phase-contrast and electron-microscopic study. Mycopathol Mycol Appl 45, 327–335

95. Dange, T., Smith, D., Noy, T., Rommel, P. C., Jurzitza, L., Cordero, R. J., Legendre, A., Finley, D., Goldberg, A. L., and Schmidt, M. (2011) Blm10 protein promotes proteasomal substrate turnover by an active gating mechanism. J Biol Chem 286, 42830–42839

96. Clarke, S. C., Dumesic, P. A., Homer, C. M., O’Donoghue, A. J., La Greca, F., Pallova, L., Majer, P., Madhani, H. D., and Craik, C. S. (2016) Integrated Activity and Genetic Profiling of Secreted Peptidases in Cryptococcus neoformans Reveals an Aspartyl Peptidase Required for Low pH Survival and Virulence. PLoS Pathog 12, e1006051

97. Baker, L. G., Specht, C. A., and Lodge, J. K. (2009) Chitinases are essential for sexual development but not vegetative growth in Cryptococcus neoformans. Eukaryot Cell 8, 1692–1705

98. Chi, A., Valencia, J. C., Hu, Z. Z., Watabe, H., Yamaguchi, H., Mangini, N. J., Huang, H., Canfield, V. A., Cheng, K. C., Yang, F., Abe, R., Yamagishi, S., Shabanowitz, J., Hearing, V. J., Wu, C., Appella, E., and Hunt, D. F. (2006) Proteomic and bioinformatic characterization of the biogenesis and function of melanosomes. J Proteome Res 5, 3135–3144

99. Almagro Armenteros, J. J., Tsirigos, K. D., Sonderby, C. K., Petersen, T. N., Winther, O., Brunak, S., von Heijne, G., and Nielsen, H. (2019) SignalP 5.0 improves signal peptide predictions using deep neural networks. Nat Biotechnol

100. Fankhauser, N., and Maser, P. (2005) Identification of GPI anchor attachment signals by a Kohonen self-organizing map. Bioinformatics 21, 1846–1852

