## Supporting information for "The structural unit of melanin in the cell wall of the fungal pathogen *Cryptococcus neoformans*"

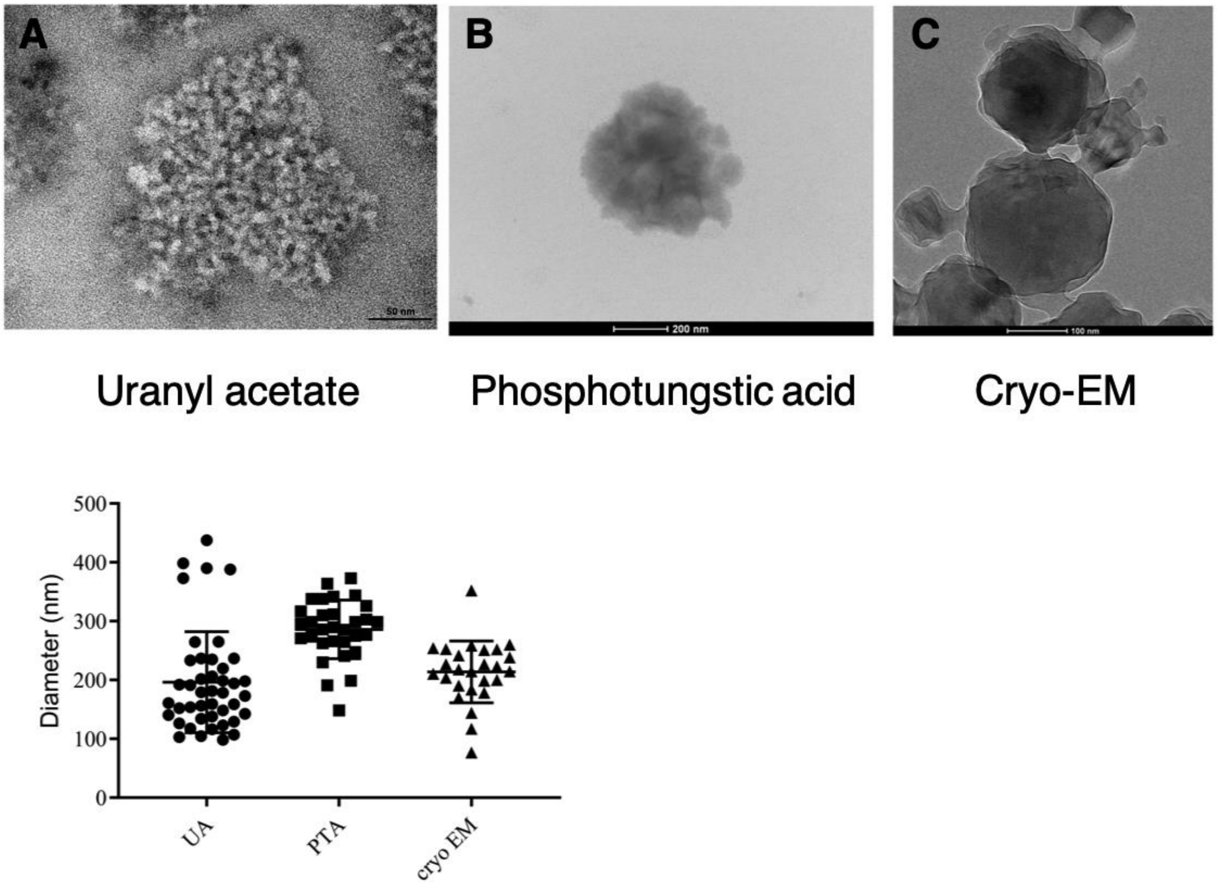

**Fig. S1 Screening and measurement of melanin granules from *C. neoformans* using different TEM approaches. A,B)** Negative staining using 2% uranyl acetate (UA) or 1% phosphotungstic acid (PTA) **C)** Cryo-EM. Melanin particles were vitreously frozen in liquid nitrogen-cooled ethane. Melanin granules showed a relatively uniform structure and comparable dimensions among all three methods. Fifty to seventy melanin granules were measured for each methodology.

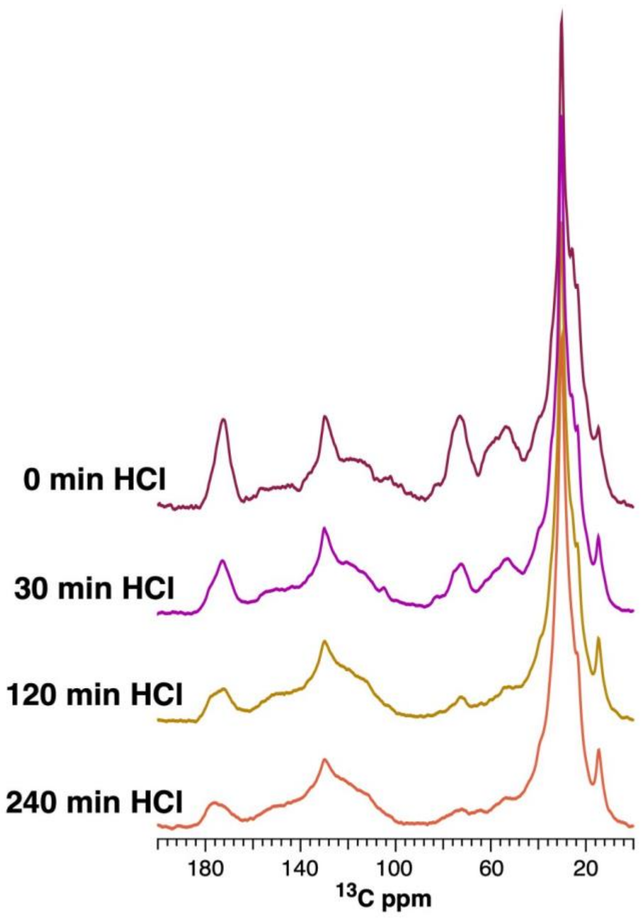

**Fig. S2 ssNMR analyses of *C. neoformans* melanin ghost after increasing incubation time with HCl demonstrate a strong association with lipids.** $^{13}C$ cross-polarization magic angle spinning (CPMAS) spectra of melanin ghosts post-acid treatment revealed that lipid signal (~30 ppm) corresponding to fatty acids is intimately associated with melanin particles.

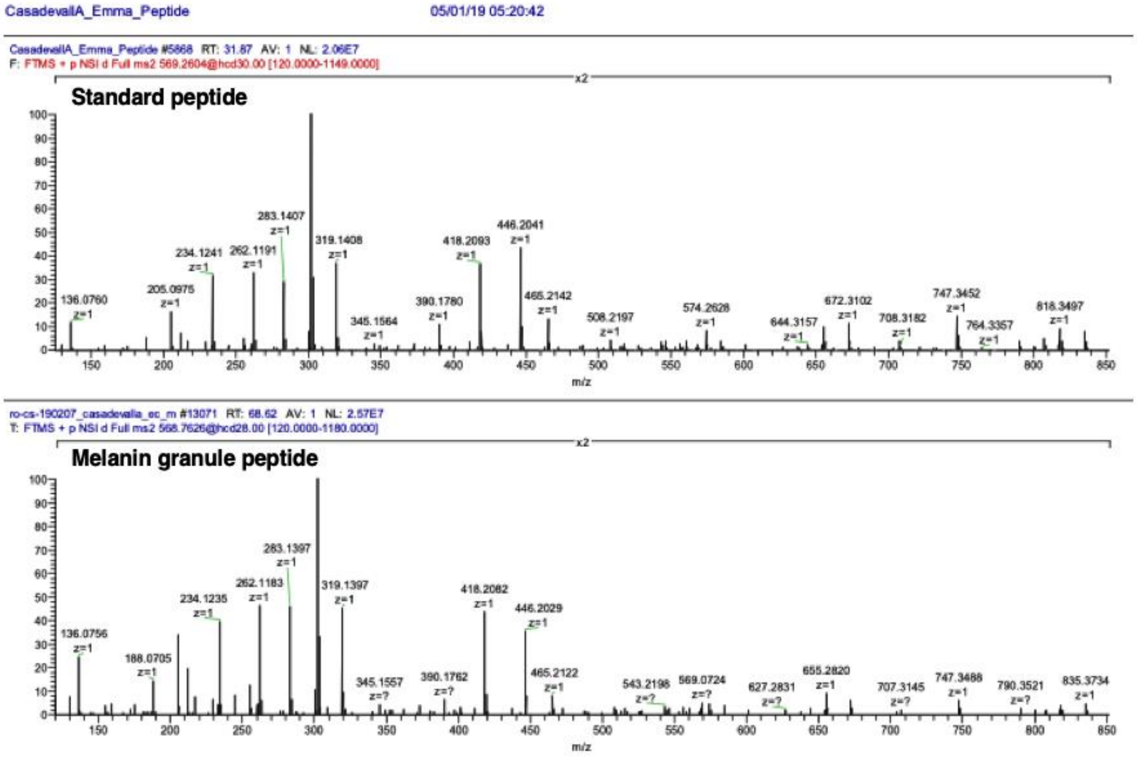

**Fig. S3 Qsp1 peptide fragmentation analysis.** Expanded view overlay of standard peptide over peptide recovered from *C. neoformans* melanin granules isolated by density gradient showing matching ions between samples thus confirming the presence of the Qsp1 peptide in the cryptococcal melanin granules.

**Table S1.** Proteins identified from *C. neoformans* melanin granules

| | Identified Proteins | Accession Number | Molecular Weight | # Unique peptide | # Unique spectra | Total spectra | Sequence coverage (%) | Peptides identified | Charge | m/z | Mass (Da) | Peptide Score | Delta PPM |
|---|---|---|---|---|---|---|---|---|---|---|---|---|---|
| 1 | Cytokine inducing-glycoprotein 1 | CNAG_01653 | 30 kDa | 18 | 38 | 238 | 29 | | | | | | |
| | | | | | | | | (R)AQNTDFETSPVAFAFPEPR(G) | 2 | 1,062.51 | 2,123.00 | 136.2 | -0.10 |
| | | | | | | | | (R)AQITDFETSPVAF(A) | 2 | 713.35 | 1,424.68 | 74.2 | 0.77 |
| | | | | | | | | (R)aQITDFETSPVAFAFPEPR(G) | 2 | 1,076.02 | 2,150.03 | 157.7 | 0.87 |
| | | | | | | | | (A)qITDFETSPVAFAFPEPR(G) | 2 | 1,048.01 | 2,094.01 | 152.0 | 1.70 |
| | | | | | | | | (T)DFETSPVAFAFPEPR(G) | 2 | 855.41 | 1,708.81 | 80.7 | 0.63 |
| | | | | | | | | (F)ETsPVAFAFPEPR(G) | 2 | 715.36 | 1,428.70 | 115.7 | -1.65 |
| | | | | | | | | (S)PVAFAFPEPR(G) | 2 | 565.80 | 1,129.59 | 67.8 | 1.40 |
| | | | | | | | | (V)AFAFPEPR(G) | 2 | 467.74 | 933.47 | 43.5 | 1.26 |
| | | | | | | | | (R)TSYPmSGGEIALVQ(Q) | 2 | 734.85 | 1,467.69 | 60.9 | -0.03 |
| | | | | | | | | (R)TSYPmSGGEIALVQQ(T) | 2 | 798.88 | 1,595.75 | 60.7 | 0.04 |
| | | | | | | | | (R)TSYPmSGGEIALVQQT(D) | 2 | 849.41 | 1,696.80 | 60.6 | 1.80 |
| | | | | | | | | (R)TSYPmSGGEIALVQQTDAQ(N) | 2 | 1,006.47 | 2,010.92 | 85.2 | 0.64 |
| | | | | | | | | (R)TSYPmSGGEIALVQQTDAQNVNIL(W) | 3 | 855.75 | 2,564.24 | 42.8 | -0.43 |
| | | | | | | | | (A)LVQQTDAQNVNILWTSESDPTR(F) | 3 | 839.09 | 2,514.24 | 101.0 | 0.72 |
| | | | | | | | | (Q)QTDAQNVNILWtSESDPTR(F) | 2 | 1,087.01 | 2,172.01 | 133.5 | -1.00 |
| | | | | | | | | (Q)NVNILWTSESDPTR(F) | 2 | 816.40 | 1,630.79 | 68.5 | -3.00 |
| | | | | | | | | (R)FHSFSTYSNSIR(E) | 3 | 482.57 | 1,444.67 | 98.8 | 0.28 |
| | | | | | | | | (R)EIGAGHYcQGAPDFSTL(G) | 2 | 988.40 | 1,974.78 | 112.9 | 0.36 |
| 2 | Quorum sensing-like peptide 1 | CNAG_03012 | 5 kDa | 1 | 1 | 6 | 29 | | | | | | |
| | | | | | | | | (N)NFGAPGGAYPW(-) | 2 | 568.76 | 1,135.51 | 57.0 | 0.60 |
| 3 | Uncharacterized protein | CNAG_05312 | 42 kDa | 1 | 1 | 6 | 3.5 | (K)VSYVQVTGVGDLTK(L) | 2 | 733.40 | 1,464.78 | 111.0 | -0.17 |
| 4 | Uncharacterized protein | CNAG_06346 | 15 kDa | 5 | 9 | 34 | 54 | (R)ATFYSPVGIGAc(G) | 2 | 741.31 | 1,480.60 | 84.5 | 0.84 |
| | | | | | | | | (R)ATFYSPVGIGACG(W) | 2 | 665.31 | 1,328.61 | 60.7 | 0.81 |
| | | | | | | | | (L)NAPQYALNADHNcGQSVR(I) | 3 | 722.98 | 2,165.90 | 115.0 | -0.15 |
| | | | | | | | | (K)VVDLCpGCNDGDLDMSPALFGAL(N) | 2 | 1168.51 | 2,335.02 | 81.0 | -0.98 |
| | | | | | | | | (L)NNNDFDQGVFPISwNFLPR(D) | 2 | 170.69 | 1,142.54 | 171.0 | -0.15 |
